# Viral microRNA regulation of Akt is necessary for reactivation of Human Cytomegalovirus from latency in CD34^+^ hematopoietic progenitor cells and humanized mice

**DOI:** 10.1101/2024.05.24.595672

**Authors:** Nicole L. Diggins, Andrew H. Pham, Jennifer Mitchell, Christopher J. Parkins, Luke Slind, Rebekah Turner, Patrizia Caposio, Jay A. Nelson, Meaghan H. Hancock

## Abstract

Human cytomegalovirus (HCMV) actively manipulates cellular signaling pathways to benefit viral replication. Phosphatidyl-inositol 3-kinase (PI3K)/Akt signaling is an important negative regulator of HCMV replication, and during lytic infection the virus utilizes pUL38 to limit Akt phosphorylation and activation. During latency, PI3K/Akt signaling also limits virus replication, but how this is overcome at the time of reactivation is unknown. Virally encoded microRNAs (miRNAs) are a key component of the virus arsenal used to alter signaling during latency and reactivation. In the present study we show that three HCMV miRNAs (miR-UL36, miR-UL112 and miR-UL148D) downregulate Akt expression and attenuate downstream signaling, resulting in the activation of FOXO3a and enhanced internal promoter-driven IE transcription. A virus lacking expression of all three miRNAs is unable to reactivate from latency both in CD34^+^ hematopoietic progenitor cells and in a humanized mouse model of HCMV infection, however downregulating Akt restores the ability of the mutant virus to replicate. These findings highlight the negative role Akt signaling plays in HCMV replication in lytic and latent infection and how the virus has evolved miRNA-mediated countermeasures to promote successful reactivation.

**AUTHOR SUMMARY:** Human cytomegalovirus (HCMV) infection results in lifelong persistence of the virus through the establishment of latency, and viral reactivation is a significant cause of morbidity and mortality in solid organ and stem cell transplant patients. HCMV latency is established in CD34^+^ hematopoietic progenitor cells (HPCs) where the virus manipulates cell signaling pathways to maintain the viral genome and remain poised to reinitiate gene expression under the appropriate conditions, although the molecular mechanisms surrounding these processes are poorly understood. HCMV encodes microRNAs (miRNAs) that modulate expression of hundreds of cellular and viral genes and play important roles in regulating signaling in HPCs. In this study, we show that HCMV miR-UL36, miR-UL112, and miR-UL148D coordinately inhibit Akt expression, activation, and downstream signaling through nonconventional mechanisms. A mutant lacking these miRNAs is unable to reactivate from latency, yet complementing Akt regulation restores the ability of the mutant virus to reactivate, pointing to an important role for miRNA-mediated inhibition of Akt to promote HCMV reactivation.

## INTRODUCTION

Human cytomegalovirus (HCMV) infects most of the world population and institutes lifelong persistence in the host through the establishment of latent infections (1). Latency is defined by maintenance of the viral genome in the absence of new virus production and occurs in CD34^+^ hematopoietic progenitor cells (HPCs) in the bone marrow and CD14^+^ monocytes (2, 3). Latent infection is punctuated by sporadic reactivation events that are stringently controlled by robust T cell responses in immunocompetent hosts. However, HCMV reactivation remains a significant cause of morbidity and mortality in the immunocompromised, including solid organ and hematopoietic stem cell transplant recipients (4, 5), and is the leading cause of viral congenital infection (6). Given its clinical importance, understanding how the virus manipulates infected HPCs during latency and reactivation is essential for developing novel approaches to target the latent reservoir.

Due to their long co-evolution with their hosts, CMVs have become master regulators of their environment, significantly remodeling cellular processes to benefit the virus lifecycle. Latency occurs in cell types inherently sensitive to intra– and extracellular cues that can drive the cells to proliferate, differentiate or undergo apoptosis. Thus, the cellular environment must be carefully controlled by viral gene products for successful latent infection and timely reactivation. Virally encoded miRNAs have emerged as important regulators of cell signaling during latency and reactivation due to their non-immunogenic nature and ability to act as rheostats, controlling signaling from external and internal stimuli to aid the virus lifecycle (7–9). Small changes to miRNA-modulated signaling pathways disrupt the careful balance necessary for latency and/or reactivation and can mediate significant phenotypic effects as evidenced by the importance of HCMV miRNA regulation of TGFβ, MAPK and RhoA signaling in CD34^+^ HPCs (10–13).

The phosphatidyl-inositol 3-kinase (PI3K)/Akt pathway is a central regulator of cell state in response to external stimuli and a common target for manipulation by viruses (14–21). Recruitment and activation of PI3K downstream of receptor tyrosine kinases, cytokine receptors, or G protein-coupled receptors promotes the conversion of phosphoinositol 4,5-bisphosphate (PIP2) to phosphoinositol 3,4,5-triphosphate (PIP3), which in turn recruits the serine/threonine kinase Akt to the membrane where it is phosphorylated at T308 by PDK1 and subsequently at S473 by mTORC2. Fully active Akt then dissociates from the membrane and phosphorylates downstream substrates to mediate changes in cell homeostasis, including enhanced protein synthesis, differentiation and regulation of stress responses (22–26). HCMV attachment and entry stimulates Akt phosphorylation, but this modification is rapidly diminished in permissive fibroblasts by pUL38 (27). The importance of diminished Akt activity in HCMV infection was shown by Zhang et al (28), who demonstrated that expression of a constitutively active Akt impairs virus replication. However, Akt signaling is necessary to stimulate protein translation and so pUL38 also directly and indirectly activates mTORC2 to bypass the need for Akt-mediated phosphorylation (29–31), highlighting how HCMV re-wires signaling pathways to benefit virus replication. Diminished Akt activity is also necessary to maintain FOXO3a nuclear localization during infection, which is normally inhibited by Akt-mediated phosphorylation (32). FOXO3a is a transcription factor that regulates differentiation and stress responses and was recently shown to be a key Akt substrate necessary for efficient HCMV replication (28), although how FOXO3a aids in virus replication remains to be defined. In the context of latency, addition of PI3K or Akt inhibitors during latency enhances HCMV replication (33), suggesting that Akt signaling is also inhibitory to virus replication in CD34^+^ HPCs. Intriguingly, FOXO3a binding sites in the HCMV major immediate early promoter are necessary for expression of immediate early genes and reactivation from latency in CD34^+^ HPCs (34), supporting the hypothesis that inactivation of Akt is a critical component of the reactivation process, although if and how this occurs has not been investigated.

Here we show that, indeed, Akt signaling is inhibitory to HCMV reactivation in CD34^+^ HPCs. Furthermore, we demonstrate that three HCMV-encoded miRNAs (miR-UL36, miR-UL112-3p and miR-UL148D-3p) regulate Akt expression that ultimately contributes to altered downstream signaling, including FOXO3a activation and expression of FOXO3a-dependent viral transcripts. Moreover, we demonstrate that regulation of Akt is one function of the three HCMV miRNAs required for efficient reactivation from latency in CD34^+^ HPC and for the first time demonstrate the importance of HCMV miRNAs in latency and reactivation *in vivo* using a humanized mouse model. These data highlight the intricate role played by Akt signaling during different aspects of the HCMV replication cycle and how viral miRNAs play an essential role in tipping the balance from latency to reactivation by modulating the outcome of Akt signaling.

## RESULTS

### Akt signaling attenuates HCMV reactivation from latency in CD34^+^ HPCs

HCMV-mediated downregulation of Akt signaling is critical for efficient lytic replication, while intact Akt signaling prevents virus replication in CD34^+^ HPCs (28, 33, 34), suggesting that Akt acts to limit virus replication in multiple cell types. At the time of reactivation, viral replication is re-initiated, but the role played by Akt and its effectors at this stage of the virus lifecycle is unknown. To address this question, we assessed the effects of Afuresertib, which inhibits the kinase activity of Akt (35, 36), and BAY1125976, which inhibits Akt phosphorylation (37) on virus reactivation in CD34^+^ HPCs. In our hands, Afuresertib modestly inhibited Akt phosphorylation but clearly disrupted downstream Akt signaling in response to EGF treatment in fibroblasts (Fig S1 A-C) while BAY1125976 inhibited phosphorylation of Akt at T308 and S473 in both HCMV-infected and uninfected fibroblasts (Fig S2 A, B). To ensure that treatment with Akt inhibitors did not adversely affect CD34^+^ HPC viability, we measured cytotoxicity in HPCs treated with each inhibitor and did not observe changes in metabolic activity at the concentrations used in these studies (Fig S1D and S2C). To examine the role of Akt signaling at the time of reactivation, human embryonic stem cell (hESC)-derived CD34^+^ HPCs were infected with TB40/E-GFP for 48 hours, and viable, CD34^+^, GFP^+^ cells were sorted and seeded into long-term bone marrow culture (LTBMC) over stromal cell support to allow for the establishment of latent infection. After 12 days of LTBMC culture, HPCs were stimulated to differentiate and re-initiate virus replication by seeding onto monolayers of permissive fibroblasts in an extreme limiting dilution assay (ELDA) in cytokine-rich media supplemented with Afuresertib, BAY1125976 or DMSO. An equivalent number of cells were mechanically lysed and plated similarly to measure free infectious virus and serve as a pre-reactivation control (38, 39) As shown in Figure 1, we observed an increase in infectious centers when cells were treated with either Afuresertib (p<0.005) or BAY1125976 (p<0.0001). The enhanced reactivation was not due to effects on HCMV replication in fibroblasts, as treatment of fibroblasts with either inhibitor does not alter HCMV replication kinetics compared to DMSO treatment (Fig S1E-H and S2 D-G). These data support the hypothesis that Akt signaling restricts HCMV reactivation in CD34^+^ HPCs.

**Figure 1.**
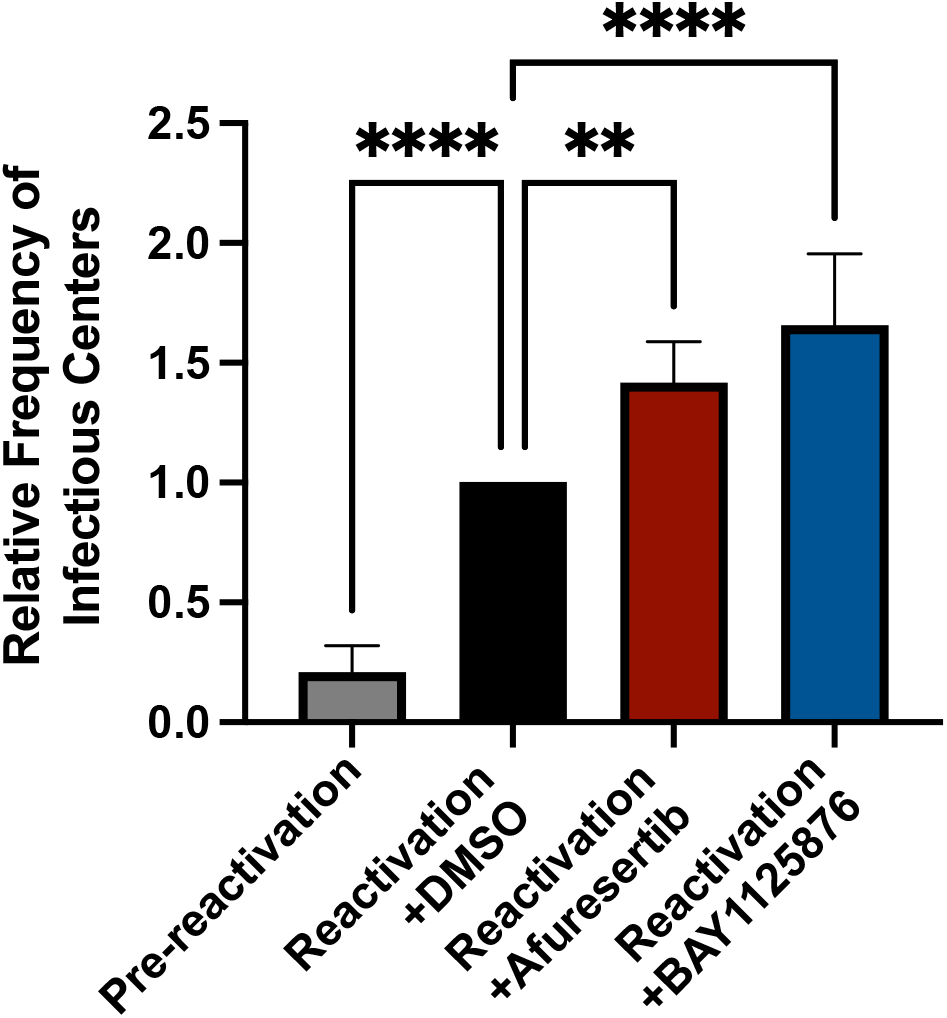
Akt restricts HCMV reactivation in CD34^+^ HPCs. hESC-derived CD34^+^ HPCs were infected with HCMV TB40/E-GFP for 48hr and then sorted by FACS for viable, CD34^+^, GFP^+^ cells. Infected HPCs were maintained in LTBMC culture medium in transwells over stromal cells for 12 days to establish latency. Latently-infected cells were co-cultured with NHDFs in cytokine-rich media in the presence of Afuresertib (100 nM), BAY1125876 (50nM), or DMSO (control) in an extreme limiting dilution assay (ELDA) to measure virus reactivation (78). An equal number of cells were mechanically disrupted and seeded in parallel to measure infectious virus present in the latency culture (pre-reactivation). At 21 days post-plating, the number of GFP^+^ wells were counted and the frequency of infectious center production was determined by ELDA software (81). Reactivation is shown as the relative frequency of infectious centers compared to DMSO control-treated cells. (**p<0.005, ****p<0.0001 [one-way ANOVA with Tukey’s multiple comparison test])

### HCMV miRNAs modulate Akt expression via multiple indirect mechanisms

Since our data indicates that Akt activity is repressive to HCMV reactivation in CD34^+^ HPCs, we hypothesized that HCMV has evolved mechanisms to inhibit Akt at this critical time in the virus lifecycle. We have previously shown that HCMV-encoded miRNAs play significant roles in CD34^+^ HPC infection (10, 11, 40) and can disrupt signaling pathways necessary for efficient reactivation from latency (12). Knowing this, we sought to determine whether any of the HCMV-encoded miRNAs affect Akt expression. To this end, we transfected HCMV or negative control miRNA mimics into normal human dermal fibroblasts (NHDFs) and assessed Akt expression levels. Western blot analysis showed that miR-UL36, miR-UL112, and miR-UL148D each reduced endogenous levels of Akt ∼50% compared to negative control miRNA (Fig. 2A, B), and co-transfection of all three miRNAs decreased Akt levels by approximately 75% (Fig. 2C, D). Interestingly, transfection of only miR-UL112 and miR-UL148D did not reduce Akt levels as efficiently as when miR-UL36 was included, suggesting that all three miRNAs contribute to maximal reduction in Akt expression. We next assessed whether these HCMV miRNAs affect Akt expression during HCMV infection. To this end, we used bacterial artificial chromosome (BAC) recombineering to generate a mutant virus lacking expression of all three HCMV miRNAs in HCMV TB40/E-GFP (ΔmiR-UL36/112/148D). We infected NHDFs with wild-type (WT) HCMV or ΔmiR-UL36/112/148D, and whole cell lysates were harvested at 48– and 96-hours post-infection (hpi). Western blot analysis demonstrated a decrease in Akt expression at 96 hpi in WT-infected cells. However, infection with ΔmiR-UL36/112/148D resulted in enhanced Akt expression compared to WT at this time point (Fig. 2 E, F). The ΔmiR-UL36/112/148D virus grew with WT kinetics (Fig. S3), suggesting that the effects on Akt expression are not due to changes in replication kinetics of the virus. Taken together, the data indicate that miR-UL36, miR-UL112, and miR-UL148D contribute to reducing Akt expression during HCMV infection.

**Figure 2.**
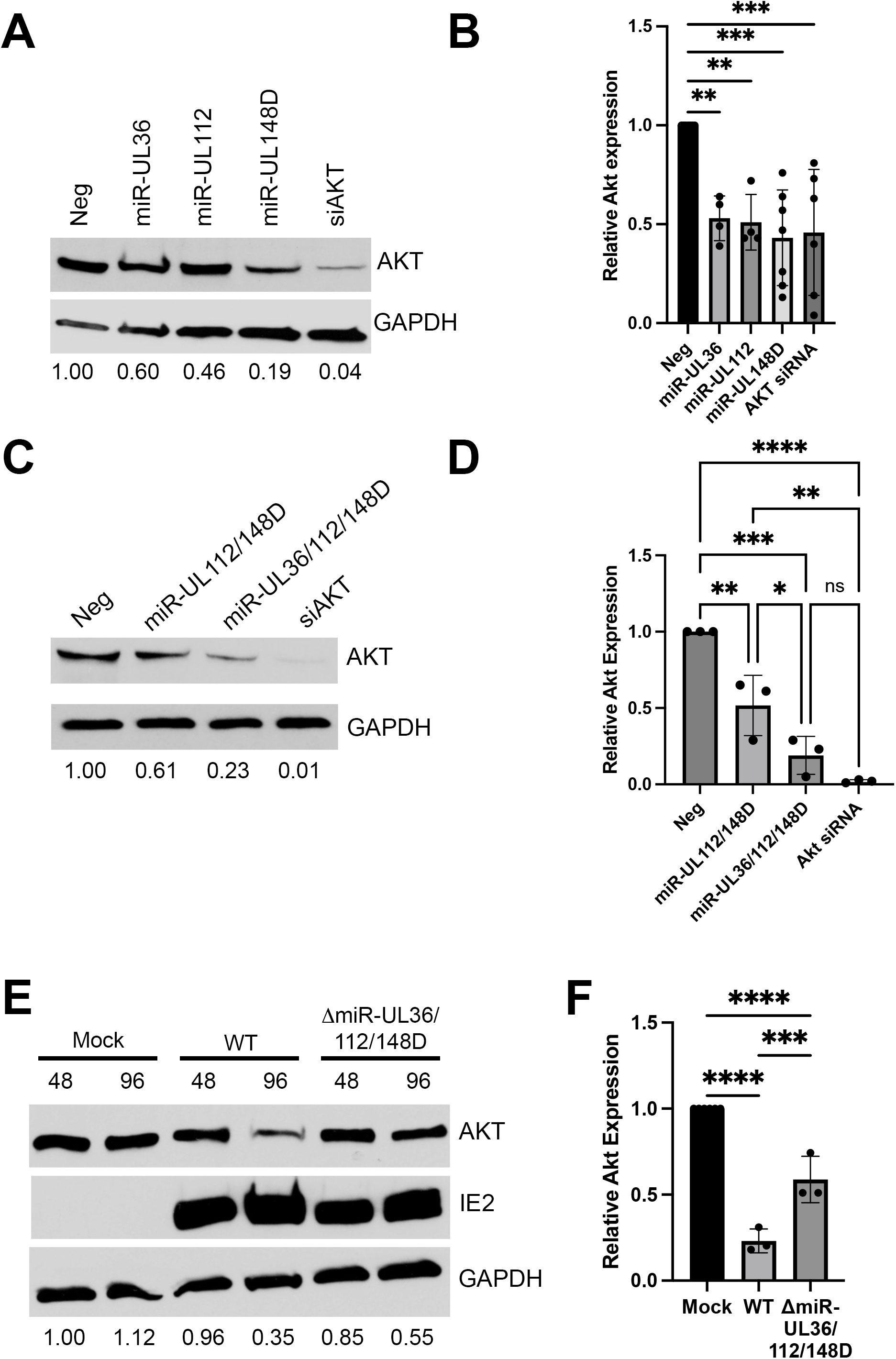
Akt is downregulated by HCMV miR-UL36, miR-UL112, and miR-UL148D. (A-D) NHDFs were transfected with double-stranded miRNA mimics, negative control (Neg), or siRNAs. Lysates were harvested 72hr post-transfection and immunoblotted for Akt and GAPDH (loading control). Quantification from one representative blot shows relative expression levels of Akt compared to Neg (normalized to GAPDH). (B, D) Quantification of (A, C), respectively, from at least three separate experiments (*p<0.05, **p<0.005, ***p<0.0005, ****p<0.0001 [one-way ANOVA]). (E) NHDFs were infected at an MOI of 3 PFU/cell with WT HCMV (TB40/E-GFP), a mutant lacking miR-UL36, miR-UL112, and miR-UL148D expression (ΔmiR-UL36/112/148D), or uninfected (Mock). Lysates were harvested 48 or 96 hpi and immunoblotted for Akt, HCMV IE2, and GAPDH. Quantification shows relative expression levels of Akt compared to Neg (normalized to GAPDH). (F) Quantification (E) at 96 hpi from six separate experiments (**p<0.005, ****p<0.0001 [one-way ANOVA with Tukey’s multiple comparison test]).

Canonical miRNA targeting occurs via complementarity between the 3’ untranslated region (UTR) of a target mRNA and the seed sequence of the miRNA (41). To test whether miR-UL36, miR-UL112, or miR-UL148D directly target the Akt 3’ UTR, we co-transfected miRNA mimics (or negative control) into HEK293T cells along with a luciferase reporter plasmid containing the 3’ UTR of Akt. To our surprise, neither miR-UL36, miR-UL112, nor miR-UL148D affected luciferase expression compared to negative control mimic (Fig. 3A), suggesting these miRNAs do not affect Akt expression by directly targeting the Akt 3’UTR. To further investigate HCMV miRNA targeting of Akt, we transfected a plasmid containing only the Akt protein coding sequence (CDS) tagged to GFP, along with HCMV miRNA mimics, Akt siRNA, or negative control mimics and whole cell lysates were harvested 24 hours post-transfection. Western blot analysis showed that miR-UL36 and miR-UL112 reduced levels of Akt-GFP compared to negative control (Fig. 3B, C), suggesting that these miRNAs affect Akt protein expression in a mechanism independent of UTR targeting. Finally, we assessed the effects of HCMV miRNAs on Akt transcript levels. NHDFs transfected with miR-UL36 or miR-UL148D, but not miR-UL112, significantly showed significantly reduced Akt transcript levels compared to negative control mimic (Fig. 3D). This suggests that miR-UL36 and miR-UL148D affect Akt protein levels by reducing mRNA levels, while miR-UL112 acts at the level of protein expression.

**Figure 3.**
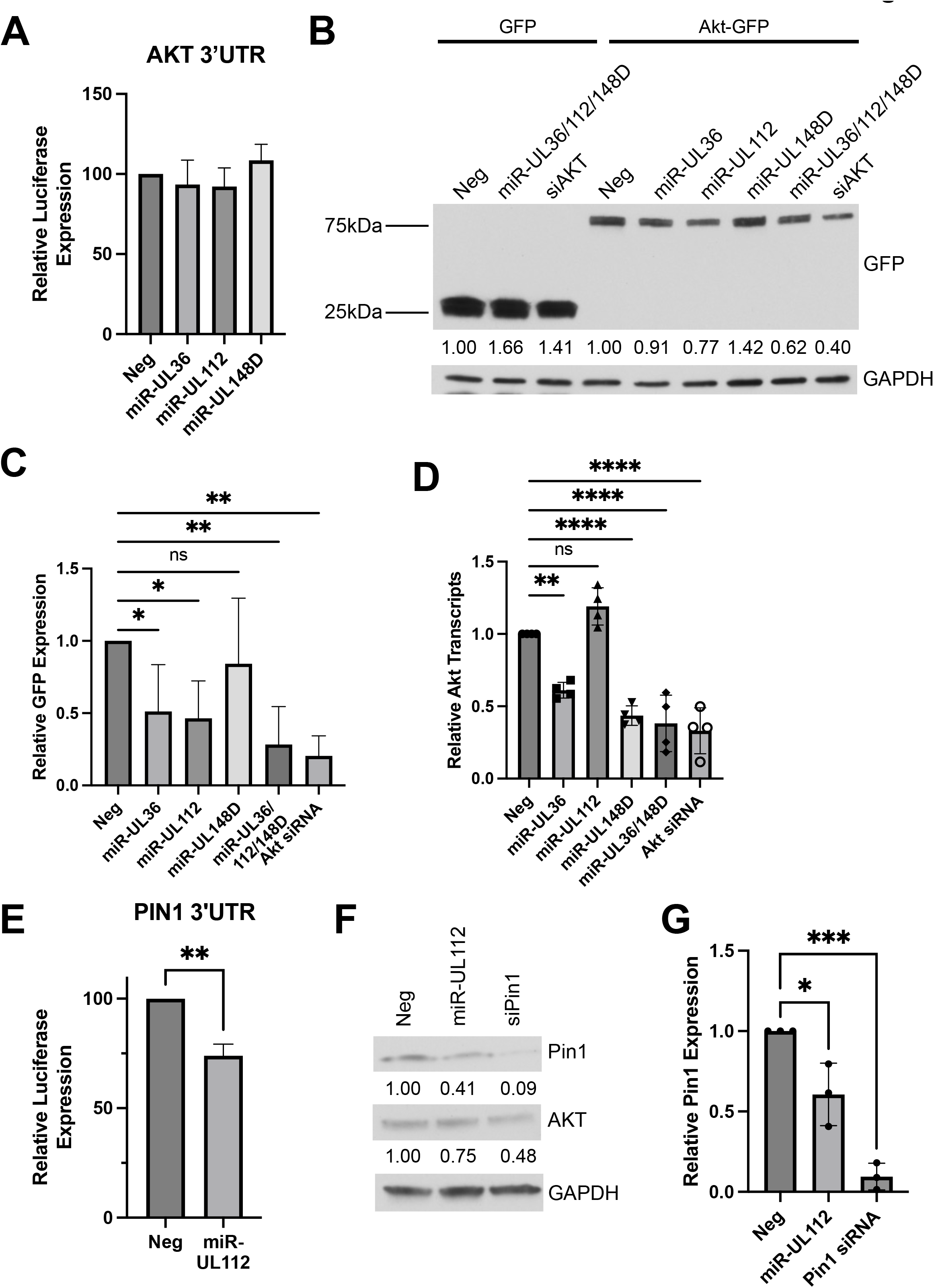
HCMV miRNAs inhibit Akt through non-canonical mechanisms. (A, E) A dual luciferase reporter containing the 3’UTR of Akt (A) or Pin1 (E) was cotransfected into HEK293T cells along with miRNA mimics. Luciferase expression was assessed 24hrs post-transfection. The relative expression is shown as a percentage of Neg. Error bars represent the standard deviation from three separate experiments (**p<0.005 by unpaired t-test). (B) HEK293T cells were co-transfected with Akt-GFP and miRNA mimics or siRNA. Lysates were harvested 24hr post-transfection and immunoblotted for GFP and GAPDH. Quantification from one representative blot shows relative expression levels of Akt-GFP compared to Neg (normalized to GAPDH). (C) Quantification of (B) from three separate experiments (*p<0.05, **p<0.005, ***p<0.0005 [one-way ANOVA with Tukey’s multiple comparison test]). (D) NHDF cells were transfected with miRNA mimics, negative control (Neg), or siRNA. RNA has harvested 72hr post-transfection, and quantitative RT-PCR for Akt was performed. Expression levels were normalized to 18S and compared to Neg (*p<0.05, **p<0.005, ***p<0.0005 [one-way ANOVA with Tukey’s multiple comparison test]). (F) NHDF were transfected with miRNA mimics or siRNA. Lysates were harvested 72hr post-transfection and immunoblotted for Pin1, Akt, and GAPDH. Quantification from one representative blot shows relative expression levels of Akt compared to Neg (normalized to GAPDH). (G) Quantification of (F) from three separate experiments (*p<0.05, ***p<0.0005 [one-way ANOVA with Tukey’s multiple comparison test]).

Since miR-UL112 reduces Akt protein expression but does not directly target the Akt 3’ UTR or affect Akt transcript levels, we hypothesized that miR-UL112 may indirectly modulate expression by targeting a regulator of Akt. One such protein is Pin1, an isomerase that promotes the stability of Akt (42). miR-UL112 decreased expression of luciferase driven by the Pin1 3’ UTR compared to negative control conditions (Fig. 3E), suggesting miR-UL112 directly targets Pin1. Moreover, transfection of miR-UL112 reduced endogenous levels of Pin1 in HEK293T cells (Fig. 3F, G) and expression of miR-UL112 or Pin1 knockdown also reduced endogenous levels of Akt (Fig. 3F), consistent with a model whereby miR-UL112 destabilizes Akt protein by reducing expression of Pin1. Taken together, our data suggest that HCMV miRNAs inhibit Akt expression via mechanisms independent of conventional 3’ UTR targeting that include affecting Akt mRNA expression and targeting regulators of Akt stability.

### miR-UL36, miR-UL112, and miR-UL148D alter Akt signaling

Knowing that Akt protein levels are reduced by HCMV miRNAs, we asked whether the miRNAs also alter signaling downstream of Akt during infection. To test this, NHDFs were infected with WT HCMV, Δ1miR-UL36/112/148D, or Mock infected for 48 hr. Cells were serum starved overnight, followed by treatment +/− EGF for 15 minutes in order to stimulate Akt phosphorylation, and lysates were collected and analyzed for phosphorylated and total protein levels. Western blot analysis showed that treatment of uninfected cells with EGF robustly induced phosphorylation of Akt at residues Threonine 308 (Fig. 4A, C) and Serine 473 (Fig. 4B, D). Cells infected with WT HCMV showed greatly reduced p-AKT levels in addition to reduced total Akt expression compared to Mock, consistent with previous reports that Akt activation is inhibited during HCMV infection (27, 28). However, cells infected with ΔmiR-UL36/112/148D consistently showed increased levels of p-Akt at both phosphorylation sites (Fig 4 A-D), along with enhanced Akt protein expression. Interestingly, we observed a striking increase in the proportion of p-Akt compared to total Akt in ΔmiR-UL36/112/148D-infected cells (Fig 4E, F), suggesting that the miRNAs alter Akt activation in addition to Akt expression. We next asked if signaling downstream of Akt is also affected by changes in total and p-Akt levels. Similar to the results with p-Akt, ΔmiR-UL36/112/148D infection resulted in enhanced phosphorylation of several Akt substrates compared to WT HCMV infection, including FOXO3a (Fig. 5A, C), PRAS40 (Fig. 5B, D), and GSK3β (Fig. S4A, C). However, compared to WT HCMV infection, no significant difference was observed in phosphorylated levels of Akt substrates mTOR (Fig. 5E, G), CREB (Fig. 5F, H), or P70S6K (Fig. S4B, D), likely as these components are direct or indirect targets of other viral gene products (29–31, 43). Together, these data demonstrate that miR-UL36, miR-UL112, and miR-UL148D reduce total and p-Akt levels, which is necessary to affect downstream AKT signaling pathways not co-opted by other virus-mediated processes.

**Figure 4.**
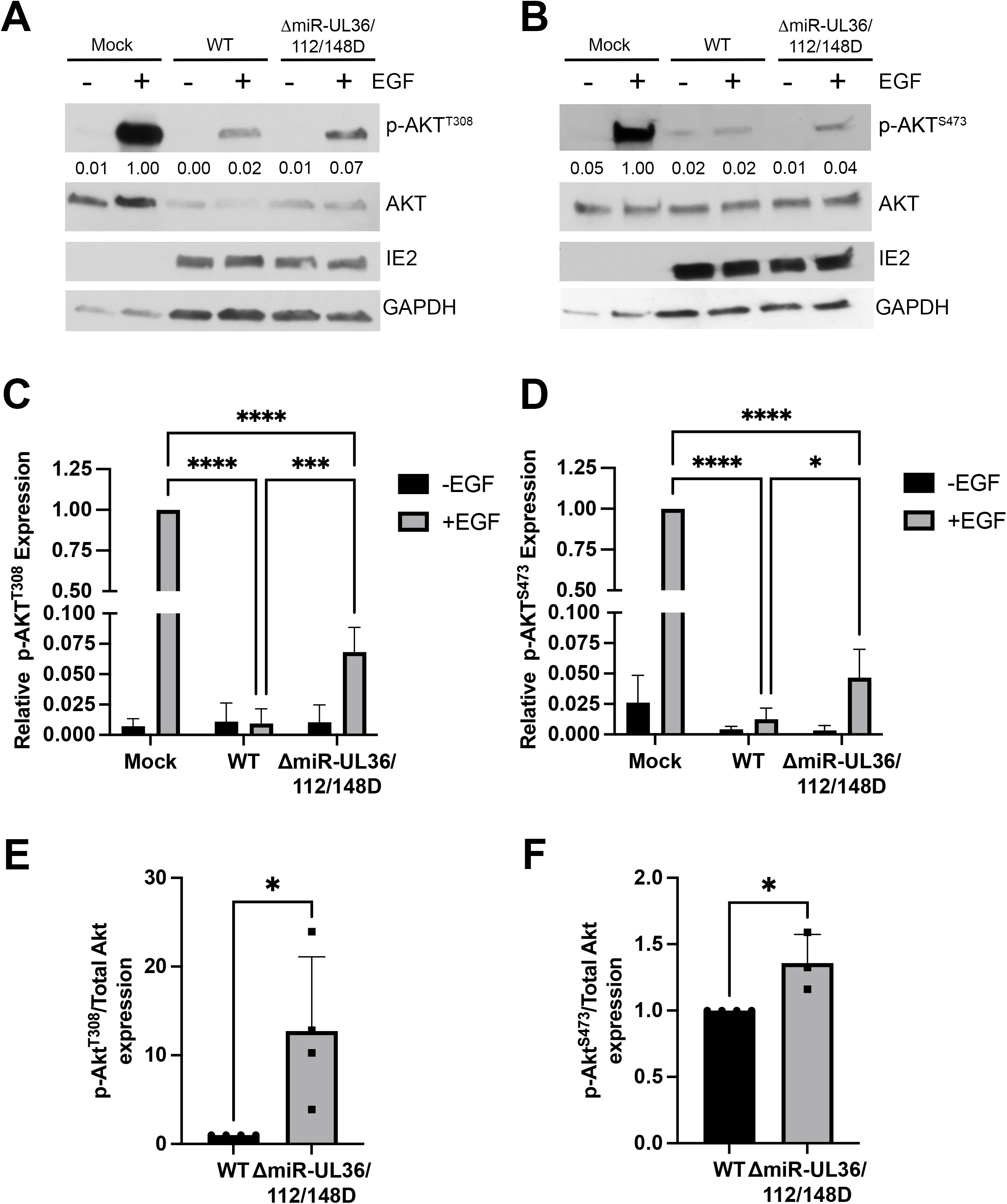
HMCV miRNAs inhibit Akt phosphorylation. (A,B) NHDF were infected with WT, ΔmiR-UL36/112/148D, or Mock infected for 48hr, serum starved overnight, and then stimulated +/−EGF for 15 minutes. Lysates were harvested and immunoblotted for Akt phosphorylated at T308 (A) or S473 (B) as well as total Akt, HCMV IE2, and GAPDH. Quantification from one representative blot shows relative expression levels of p-Akt compared to Neg (normalized to GAPDH). (C, D) Quantification of p-Akt levels from (A, B), respectively, from three separate experiments (comparing +EGF conditions *p<0.05, ***p<0.0005, ****p<0.0001 [two-way ANOVA with Tukey’s multiple comparison test]). (E, F) Ratio of p-Akt to total Akt levels in WT and ΔmiR-UL36/112/148D from (A, B), respectively, from three separate experiments, normalized to WT (*p<0.05 [unpaired t-test]).

**Figure 5.**
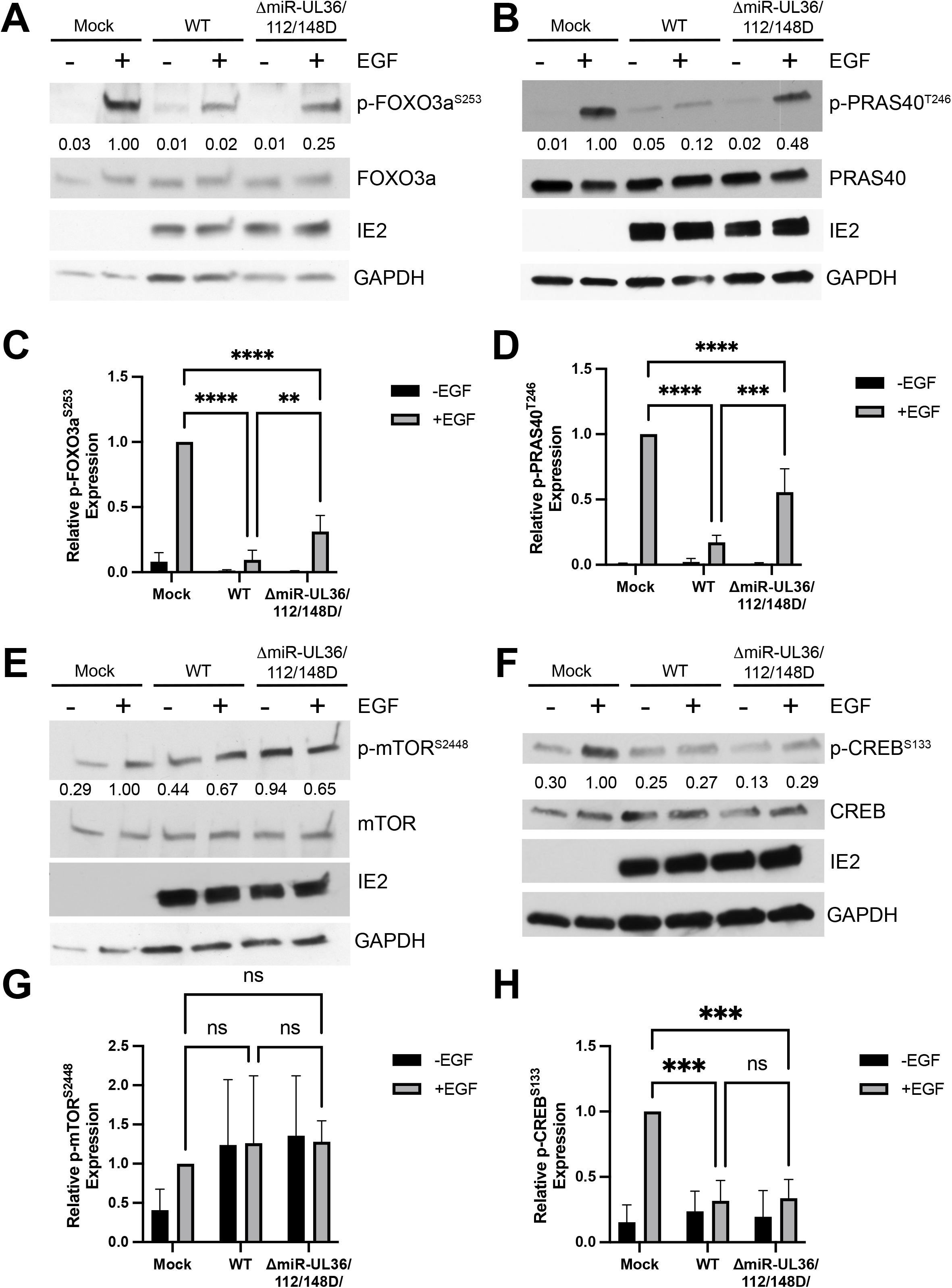
HCMV miRNAs affect signaling downstream of Akt. (A, B, E, F) NHDF were infected with WT, ΔmiR-UL36/112/148D, or Mock infected for 48hr, serum starved overnight, and then stimulated +/−EGF for 15 minutes. Lysates were harvested and immunoblotted for indicated phosphorylated and total proteins as well as HCMV IE2 and GAPDH. Quantification from one representative blot shows relative expression levels of p-protein compared to Neg (normalized to GAPDH). (C, D, G, H) Quantification of (A, B, E, F), respectively, from three separate experiments (comparing +EGF conditions, **p<0.005, ***p<0.0005, ****p<0.0001 [two-way ANOVA with Tukey’s multiple comparison test]).

### HCMV miRNAs affect FOXO3a nuclear localization and function

FOXO3a is an important effector regulated by Akt signaling during HCMV infection (34, 44). FOXO3a in its active, unphosphorylated form localizes to the nucleus and acts as a transcription factor to regulate cellular homeostasis, stress responses, and apoptosis (32, 45). Our data suggest that HCMV miR-UL36, miR-UL112, and miR-UL148D together reduce FOXO3a phosphorylation (Fig 5A-C). Thus, we hypothesized that HCMV miRNAs promote nuclear translocation of FOXO3a. To test this, NHDFs were infected with WT HCMV, ΔmiR-UL36/112/148D, or Mock infected for 72hr and immunostained for FOXO3a, DAPI, and actin (phalloidin). Cells infected with WT HCMV showed an ∼64% increase in FOXO3a nuclear localization compared to mock (Figs. 6A, B), consistent with unphosphorylated, active FOXO3a in WT-infected cells. In cells infected with the ΔmiR-UL36/112/148D mutant, FOXO3a localized throughout the cell (Fig. 6A) and nuclear FOXO3a levels were not significantly different from mock infection (Fig. 6B).

**Figure 6.**
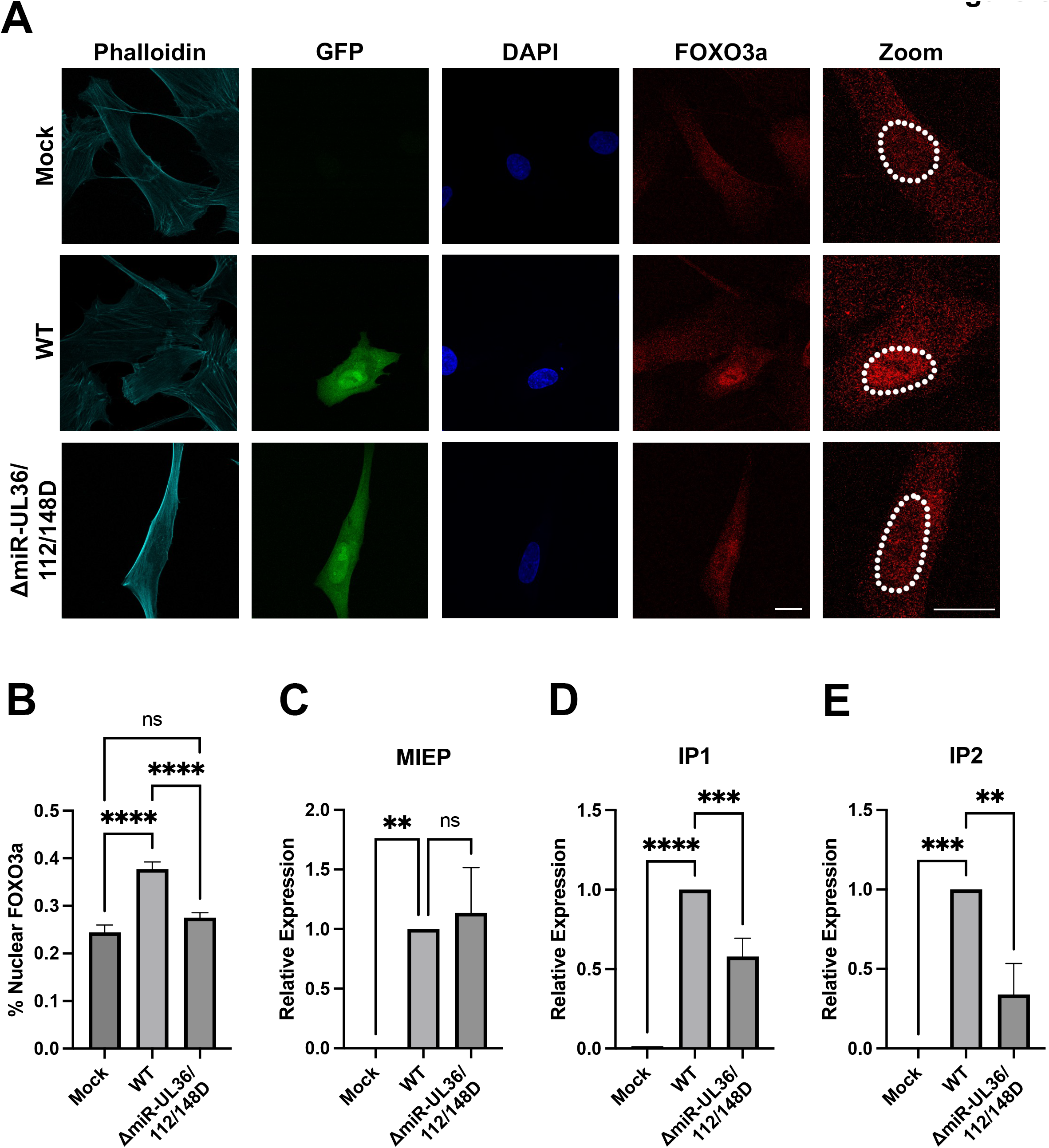
HCMV miRNAs promote FOXO3a nuclear localization and induction of MIE transcripts. (A, B) NHDF were plated on coverslips and infected with WT, ΔmiR-UL36/112/148D, or Mock infected. Cells were fixed 72 hpi and stained for actin (phalloidin), FOXO3a, and nuclei (DAPI). (A) Representative images are shown. White dotted line shows outline of the nucleus in FOXO3a images. Scale bar, 20μM. (B) Image J software was used to quantify the average intensity of FOXO3a in the nucleus and the entire cell and graphed as a percentage of nuclear FOXO3a levels normalized to the whole cell. Error bars represent the standard error of the mean for 38-44 cells from each condition from three separate experiments (****p<0.0001 [one-way ANOVA with Tukey’s multiple comparison test]). (C-E) NHDF were infected with WT, ΔmiR-UL36/112/148D, or Mock infected for 72hr and RNA has harvested. Quantitative RT-PCR was performed using specific primers for MEIP (C), IP1 (D), or IP2 (E). Expression levels were normalized to GAPDH and compared to Mock (**p<0.003, ***p<0.0007, ****p<0.0001 [one-way ANOVA with Tukey’s multiple comparison test]).

Recent work has identified two alternative intronic promoters containing FOXO3a binding sites (iP1 and iP2) in the major immediate early (MIE) locus that play an important role in stimulating IE gene expression during reactivation from latency in CD34^+^ HPCs (34, 46). Given our observations that HCMV miR-UL36, miR-UL112, and miR-UL148D promote FOXO3a activation during HCMV infection (Fig. 5 A, C and Fig. 6 A, B), we hypothesized that a downstream consequence would be the induction of iP1 and iP2 transcripts. To test this, we infected NHDF with WT HCMV, ΔmiR-UL36/112/148D, or mock infected, harvested RNA 72 hours later and performed qPCR for MIEP-, iP1-, or iP2-derived transcripts. While infection with ΔmiR-UL36/112/148D showed similar levels of MIEP transcripts to WT infection (Fig. 6C), ΔmiR-UL36/112/148D-infected cells produced significantly lower amounts (p=0.0006 and p=0.0022, respectively) of transcripts derived from iP1 and iP2 promoters (Fig. 6D, E, respectively). Together, these data suggest that inhibition of Akt by HCMV miRNAs results in reduced FOXO3a phosphorylation, increased translocation to the nucleus, and IE gene transcription from promoters that are important for reactivation from latency.

### HCMV miR-UL36, miR-UL112, and miR-UL148D promote reactivation from latency *in vitro* **and *in vivo***

Since we have shown that Akt signaling impairs virus reactivation (Fig. 1) and miR-UL36, miR-UL112, and miR-UL148D reduce Akt protein levels (Fig. 2, 3, 4) and IE transcripts from promoters important for reactivation (Fig. 6D, E), we next assessed the ability of the ΔmiR-UL36/112/148D mutant to reactivate from latency in CD34^+^ HPCs. Infection of hESC-derived CD34^+^ HPCs with the ΔmiR-UL36/112/148D mutant infected cells showed a significant decrease (p<0.0001) in the frequency of infectious center production compared to WT infected cells (Fig. 7A). Moreover, the frequency of reactivation in ΔmiR-UL36/112/148D-infected cells was not significantly different than the pre-reactivation control, suggesting that when HCMV is lacking miR-UL36, miR-UL112, and miR-UL148D, the virus is unable to reactivate from latency efficiently. HCMV genomes levels were similar for both viruses at the beginning and end of latency (Fig. 7B), indicating that the reactivation defect of the ΔmiR-UL36/112/148D mutant is not due to differences in initial binding and entry or a loss of viral genomes or genome-containing cells during latency.

**Figure 7.**
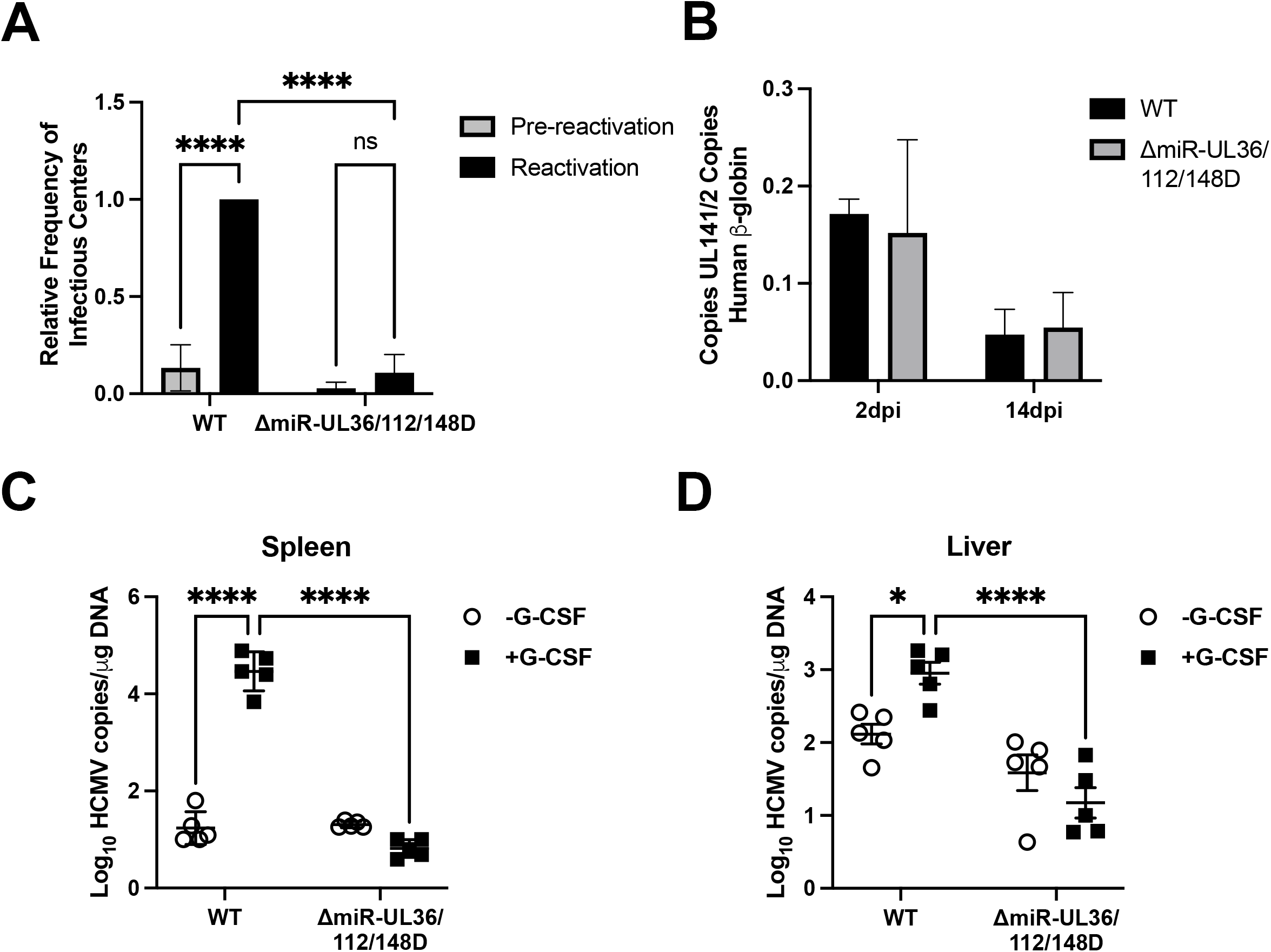
miR-UL36, miR-UL112, and miR-UL148D are important for HCMV reactivation from latency *in vitro* and *in vivo*. (A, B) hESC-derived CD34^+^ HPCs were infected with WT, ΔmiR-UL36/112/148D, or Mock for 48hr and then sorted by FACS for viable, CD34^+^, GFP^+^ cells. Infected HPCs were maintained in LTBMC culture medium in transwells over stromal cells for 12 days to establish latency. (A) Latently-infected cells were co-cultured with NHDFs in cytokine-rich media in an extreme limiting dilution assay (ELDA) to measure viral reactivation. An equal number of cells were mechanically disrupted and seeded in parallel to measure infectious virus present in the latency culture (pre-reactivation). At 21 days post-plating, the number of GFP^+^ wells were scored and the frequency of infectious center production was determined by ELDA software. Reactivation is shown as the relative frequency of infectious center compared to WT infected cells. (B) Total genomic DNA was isolated from HPCs at 2 days post infection (dpi) (post-sort) or 14 dpi (latency day 12), and quantitative real-time PCR was used to quantify the ratio of viral genomes (copies of HCMV UL141) to cellular genomes (per two copies of human β-globin). Error bars represent standard deviation from triplicate samples. Data shown are representative of three independent experiments. (C, D) Sub-lethally irradiated NOD-*scid* IL2Rψ ^null^ mice were engrafted with CD34^+^ HPCs (huNSG) and subsequently injected with human fibroblasts previously infected with HCMV WT or ΔmiR-UL36/112/148D. At 4 weeks post-infection, viral reactivation was triggered by treating latently infect HCMV WT and ΔmiR-UL36/112/148D (n=5) with G-CSF and AMD-3100. At 1 week post-treatment, mice were euthanized, and tissues were harvested. Total genomic DNA was isolated from spleen tissue (C) or liver tissue (D), and HCMV genomes were quantified using qPCR with primers and probes specific for the UL141 gene (*p<0.05, ***p<0.0005, ****p<0.0001 [two-way ANOVA followed by Bonferroni’s multiple comparison test]).

To further support the findings that a ΔmiR-UL36/112/148D virus is unable to reactivate from latency, we assessed reactivation of this virus in an *in vivo* humanized mouse model (47, 48). CD34^+^ HPCs were engrafted into NOD-*scid*IL2Rψ ^null^ mice (huNSG) followed by infection with WT HCMV or ΔmiR-UL36/112/148D. After viral latency was established, HCMV reactivation was stimulated by treatment of mice with granulocyte colony-stimulating factor (G-CSF) and AMD3100 which results in virus dissemination into tissues through infected macrophages (49). WT-infected animals showed significantly increased DNA copy numbers in both spleen (Fig. 7C) (p<0.0001) and liver (Fig. 7D) (p=0.031) tissues of huNSG mice following reactivation stimulus. Critically, no change in DNA copy number was observed in ΔmiR-UL36/112/148D-infected mice triggered to reactivate, demonstrating that miR-UL36, miR-UL112, and miR-UL148D are also important for reactivation from latency *in vivo*.

### Akt inhibition by HCMV miRNAs contributes to reactivation from latency

While we have shown that miR-UL36, miR-UL112-3p and miR-UL148D-3p each independently regulate Akt expression (Fig. 2), each miRNA targets several additional proteins (8, 50–61), many of which have not been investigated in the context of latency. In order to determine if the reactivation defect of the ΔmiR-UL36/112/148D mutant is due to its inability to regulate Akt protein levels specifically, we generated a mutant HCMV lacking expression of miR-UL36, miR-UL112, and miR-UL148D as well as expressing an shRNA targeting Akt in the 3’UTR of HCMV UL22A (ΔmiR-UL36/112/148D/Akt shRNA), a highly abundant transcript expressed during latency. As a control, we generated a virus expressing a non-targeting *Caenorhabditis elegans* miRNA, cel-miR-67, in this same region in both WT HCMV (WT/cel-miR-67) and the ΔmiR-UL36/112/148D mutant (ΔmiR-UL36/112/148D/cel-miR-67) (Fig. S5A). To confirm that these viruses modulate Akt levels we quantified Akt transcripts in NHDFs infected with WT/cel-miR-67, ΔmiR-UL36/112/148D/cel-miR-67, ΔmiR-UL36/112/148D/Akt shRNA, or Mock infected. Akt transcript levels were decreased in WT/cel-miR-67-infected cells at 96hpi compared to mock. NHDFs infected with ΔmiR-UL36/112/148D/cel-miR-67 showed significantly higher (p=0.044) Akt transcript levels than WT, consistent with the effects of ΔmiR-UL36/112/148D virus on Akt protein levels. As predicted, the ΔmiR-UL36/112/148D/Akt shRNA virus showed decreased Akt transcripts compared to the ΔmiR-UL36/112/148D/cel-miR-67 virus (Fig. S5B), effectively complementing the lack of Akt regulation of the ΔmiR-UL36/112/148D/cel-miR-67 mutant. As expected, neither ΔmiR-UL36/112/148D/cel-miR-67 nor ΔmiR-UL36/112/148D/Akt shRNA exhibited a significant lytic replication defect compared to WT/cel-miR-67 (Fig. S5C-F).

We next assessed the ability of the ΔmiR-UL36/112/148D/Akt shRNA virus to reactivate from latency. Like WT HCMV, the WT/cel-miR-67 virus was able to establish latency and reactivate, as indicated by this increase in the frequency of infectious centers compared to the pre-reactivation controls (Fig. 8A). Furthermore, the ΔmiR-UL36/112/148D/cel-miR-67 virus exhibited a reactivation defect (Fig. 8A) similar to the ΔmiR-UL36/112/148D virus (Fig. 7A). Moreover, genomes were maintained in ΔmiR-UL36/112/148D/cel-miR-67-infected cells compared to infection with WT/cel-miR-67 (Fig. 8B), similar to infection with ΔmiR-UL36/112/148D (Fig. 7B). Finally, infection with the ΔmiR-UL36/112/148D/Akt shRNA virus demonstrated an enhanced frequency of infectious centers compared to ΔmiR-UL36/112/148D/cel-miR-67 (Fig. 8A), but no change in genome copy number (Fig. 8B), suggesting that inhibiting Akt in the context of infection with the ΔmiR-UL36/112/148D mutant is able to partially complement the reactivation defect observed when these miRNAs are lacking during infection. Taken together, these data suggest that miR-UL36, miR-UL112, and miR-UL148D promote HCMV reactivation from latency via a mechanism at least partially dependent on reducing Akt expression.

**Figure 8.**
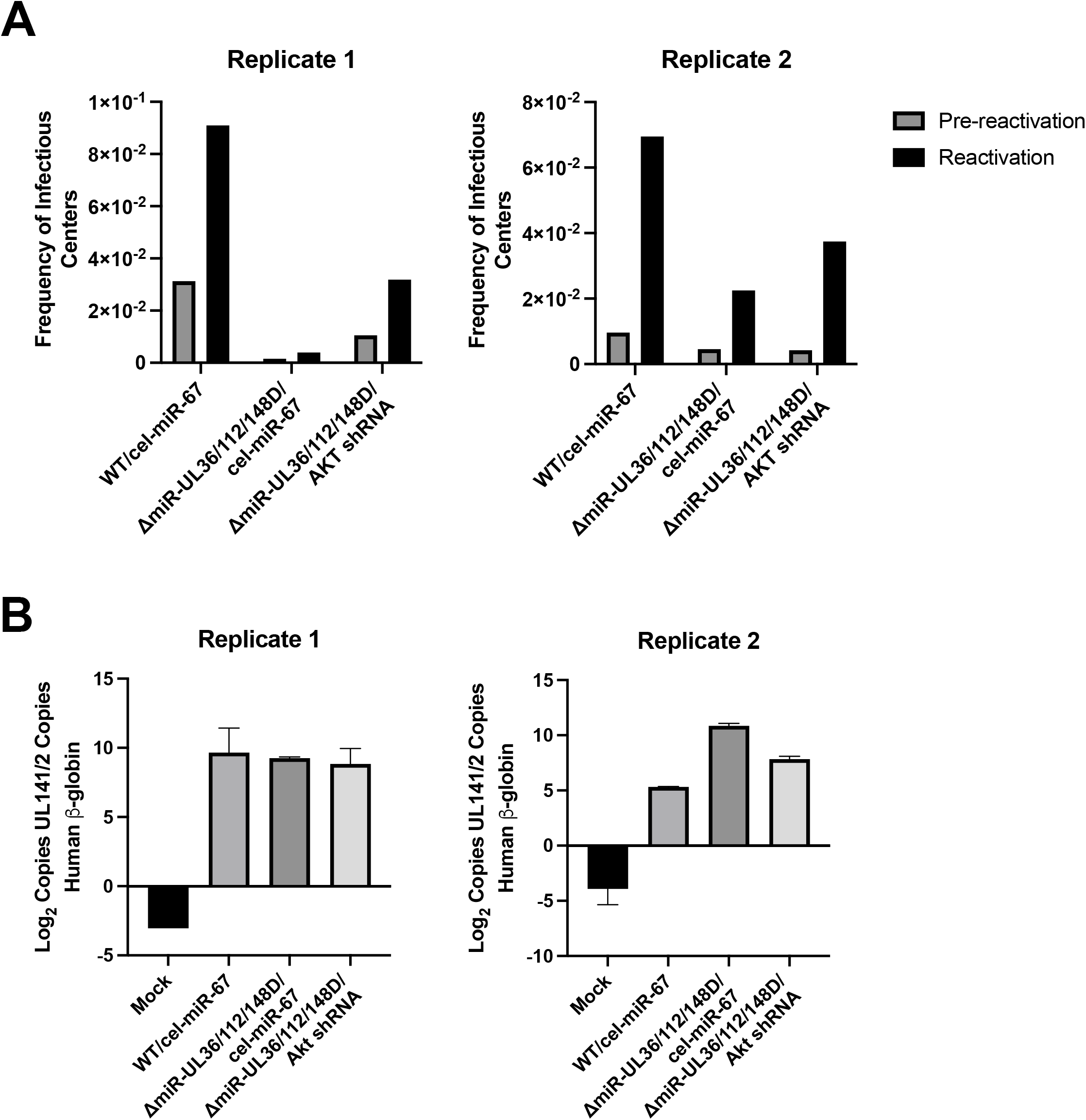
HCMV miRNA regulation of Akt contributes to reactivation from latency. (A) hESC-derived CD34^+^ HPCs were infected with WT HCMV expressing cel-miR-67 (WT/cel-miR-67), ΔmiR-UL36/112/148D/cel-miR-67, UL36/112/148D/Akt shRNA, or Mock infected for 48hr and then sorted by FACS for viable, CD34^+^, GFP^+^ cells and used in latency and reactivation assays as described in Figure 7. Results showing frequency of infectious centers from two independent experiments are shown. (B) Total genomic DNA was isolated from HPCs 14 dpi, and quantitative real-time PCR was used to quantify the ratio of viral genomes as described in Figure 7. Error bars represent standard deviation from triplicate samples. Data shown are from two representative experiments.

## DISCUSSION

In this study, we provide the first mechanistic evidence of how HCMV regulates Akt signaling at the time of reactivation from latency. We show that three miRNAs encoded by HCMV—miR-UL36, miR-UL112, and miR-UL148D—coordinately inhibit Akt expression and alter downstream Akt signaling during infection. By modulating Akt protein levels, these HCMV miRNAs prevent the phosphorylation and inactivation of FOXO3a, thereby promoting nuclear localization and inducing expression of MIE transcripts containing FOXO3a binding sites. Importantly, we show that an HCMV mutant lacking expression of the three miRNAs fails to reactivate from latency both *in vitro* and *in vivo*, yet expression of an Akt shRNA can partially complement the reactivation defect, indicating that reducing Akt expression is one mechanism used by HCMV miRNAs to promote reactivation.

Our data clearly demonstrate that miR-UL36, miR-UL112, and miR-UL148D affect Akt protein levels both when ectopically expressed (Fig 2 A-D) and during HCMV infection (Fig 2 E, F). Surprisingly, none of these miRNAs target the 3’ UTR of Akt in a canonical manner (Fig. 3A). Each miRNA reduces Akt levels independently, and through different mechanisms. miR-UL36 reduces both RNA (Fig. 3D) and protein levels (Fig. 2 A-D) and affects expression of Akt-GFP from a plasmid lacking the 3’ and 5’ UTRs (Fig. 3B, C), suggesting miR-UL36 targets the CDS of Akt. Binding of miRNAs to the CDS commonly occurs with extensive base pairing at the 3’ end of the mature miRNA (62). Indeed, we observed a region of complementarity in the Akt CDS with miR-UL36 (data not shown). However, CDS targeting typically results in translational repression without affecting mRNA levels (63, 64), although this is not always the case (62, 65). Therefore, while it is possible that miR-UL36 indirectly affects Akt expression, our data provide evidence that miR-UL36 targets the coding region of Akt. miR-UL148D, on the other hand, reduces Akt transcript (Fig. 3D) and protein levels (Fig. 2A-D), but does not target the 3’UTR (Fig. 3A) or the coding region (Fig. 3B, C). We also did not find any predicted target sites for miR-UL148D in the 5’ UTR of Akt (data not shown). Taken together, these data point to a role for miR-UL148D indirectly affecting Akt mRNA expression. The IER5 transcription factor was identified as a miR-UL148D target (53), but further study is needed to determine if transcription factor targeting by miR-UL148D is responsible for the effects on Akt transcription. Lastly, miR-UL112 expression decreases Akt protein (Fig. 2A-D) but not RNA levels (Fig. 3D) and does not target the 3’ UTR (Fig. 3A). Here we identify the isomerase Pin1 as a target of miR-UL112. Pin1 aids in stabilizing active Akt and preventing its degradation (42). Our data show that miR-UL112 targets the 3’ UTR of Pin1 (Fig. 3E) and ectopic expression of miR-UL112 decreases both Pin1 and Akt levels (Fig. 3F, G). Thus, our data suggest that one mechanism by which HCMV miRNAs reduce Akt levels is by targeting Pin1 to induce Akt degradation. These findings highlight the myriad ways that viral miRNAs can affect expression of a single protein to elicit a phenotypic effect.

Akt phosphorylation is induced upon HCMV entry into fibroblasts but is downregulated within 12 hrs post-infection (27, 28). Consistent with this, we observed decreased Akt phosphorylation in response to EGF stimulation during infection with WT HCMV. However, during infection with the Δ1miR-UL36/112/148D mutant, total and p-Akt levels were higher than during WT infection (Fig. 4), suggesting that these miRNAs act to reduce activation as well as expression of Akt. Since removing these miRNAs from HCMV only partially restored p-Akt levels compared to Mock infection, this suggests that other HCMV factors inhibit Akt activation during HCMV infection.

Indeed, HCMV UL38 contributes to inhibition of Akt phosphorylation via a negative feedback loop involving mTORC1 and IRS1 (27). Given that infection with the ΔmiR-UL36/112/148D mutant shows increased p-Akt levels despite UL38 expression suggests that these miRNAs also play a direct role in regulating Akt signaling apart from regulating total Akt levels (Fig. 4E-F). Interestingly, we observed a greater increase in phosphorylation at T308 than S473 (Fig. 4E, F) which suggests that HCMV miRNAs preferentially regulate pathways leading to T308 phosphorylation mediated by PDK1. We did not observe any changes in expression of PDK1 (data not shown), a kinase known to phosphorylate Akt at Thr308 (66), or phosphorylation of mTOR (Fig. 5E), which is part of the mTORC2 complex that phosphorylates Akt at Ser473 (67) and thus further study is needed to identify other potential regulators of Akt phosphorylation targeted by the miRNAs. Intriguingly, miR-UL36, miR-UL112, and miR-UL148D only affect activation of a subset of the tested Akt effectors, including FOXO3a, PRAS40, GSK3β, and Akt itself, but not other effectors like CREB, mTOR, and P70S6K. UL38 bypasses the need for Akt activity to maintain protein translation, making mTOR and p70S6K immune to the downstream effects of miRNA-mediated Akt regulation. HCMV regulation of CREB phosphorylation is less well understood, but CREB binding sites in the MIEP are necessary for efficient reactivation from latency (68). Our data indicate that CREB phosphorylation is regulated by a mechanism that is protected from miRNA-regulated, Akt-mediated phosphorylation (Figs. 5 and S4). Regulation of Akt via three HCMV miRNAs is also necessary to modulate FOXO3a activity and localization (Figs. 5 and 6). Interestingly, FOXO3a is a target of miR-US5-1 and miR-UL112, whose downregulation is important for inhibiting apoptosis early after infection of CD34^+^ HPCs (40). We did not observe a change in total FOXO3a expression during infection with ΔmiR-UL36/112/148D (Fig. 5A), suggesting miR-UL112 alone is insufficient to functionally affect FOXO3 levels during infection. However, our current study suggests that miR-US5-1 and miR-UL112 downregulation of FOXO3a, along with the effectors of Akt signaling, work together to block virus replication during latency establishment. Taken together, our current study, along with previously published work, highlights the growing evidence that HCMV miRNAs can target multiple components of a signaling pathway to alter downstream functional outcomes (11, 55, 59, 60, 69). Clearly, HCMV regulates Akt activation and signaling in myriad ways, and our data points to HCMV miRNAs as contributors to this regulation, most appreciably during reactivation from latency in CD34^+^ HPCs.

In CD34^+^ HPCs, EGFR/Akt signaling is required for establishment and/or maintenance of latency, as treatment of HPCs with Akt inhibitors results in enhanced virus replication compared to untreated conditions (33). We hypothesize that the intensity of Akt signaling acts as a switch between latency and reactivation; EGFR/Akt signaling is involved in reducing virus replication during latency establishment via an unknown mechanism, but Akt signaling must be attenuated at the time of reactivation in order to stimulate replication. In support of this model, treatment of HPCs with two different Akt inhibitors when cells are stimulated to reactivate results in enhanced reactivation in WT-infected cells (Fig. 1), suggesting that Akt activity restricts some aspects of the reactivation process. miRNA regulation of Akt levels contributes to the process of reactivation, and this is dependent on expression of multiple HCMV miRNAs. While miR-UL112 and miR-UL148D are expressed during latency, miR-UL36 is not detected (12). Furthermore, expression of miR-UL112 and miR-UL148D only partially reduce Akt levels compared to expression of all three miRNAs (Fig 2C and D). Therefore, we hypothesize that during latency miR-UL112 and miR-UL148D are unable to influence Akt activity enough to tip the balance towards reactivation. However, the additional expression of miR-UL36 during the early stages of reactivation reduces Akt signaling enough to have a phenotypic effect on signaling and to stimulate virus replication (Fig. 9). In agreement with this, infection with the ΔmiR-UL36/112/148D mutant, which results in increased Akt activity compared to WT-infected cells, exhibits a reactivation defect *in vitro* and *in vivo* (Fig. 7), but this defect can be partially overcome by expression of an Akt shRNA (Fig. 8). Importantly, these findings are the first to show that HCMV miRNAs have a phenotypic function *in vivo*, further underscoring the importance of virally encoded miRNAs to the HCMV lifecycle.

**Figure 9.**
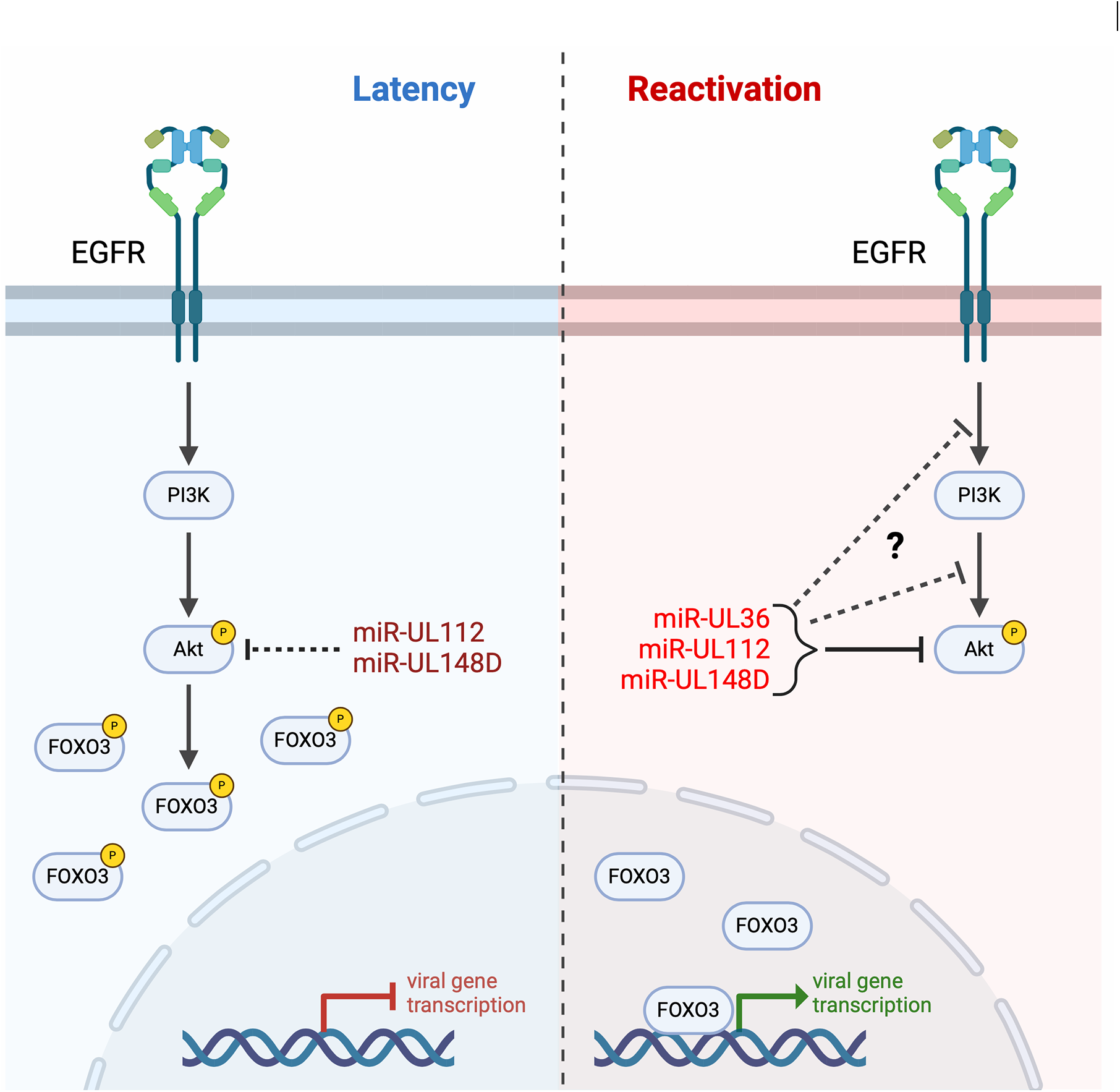
Model of Akt regulation in HCMV latency and reactivation. Left, EGFR and Akt signaling is required for latency. Active Akt promotes phosphorylation and inhibition of FOXO3a, thereby limiting viral gene transcription. Right, during reactivation, miR-UL36, miR-UL112, and miR-UL148D are expressed, which reduce Akt expression and activation. These miRNAs in turn promote active FOXO3a nuclear translocation and transcription of MIE genes.

Our data also highlight the complexity of miRNA regulation of the processes of latency and reactivation. Previous work shows that a miR-UL148D mutant virus has a replicative phenotype in CD34^+^ HPCs (53) while a miR-UL112 mutant showed no specific defect in latency or reactivation in CD14^+^ monocytes (70). When mutations to miR-UL112, miR-UL148D and miR-UL36 are combined, the resulting mutant is unable to reactivate, indicating that the replicative phenotype of the miR-UL148D mutant can be overcome with the loss of additional miRNAs. Unravelling the important function(s) of each miRNA will help to understand the phenotypes of each individual and combination mutant. Together, our data for the first time describes a miRNA-mediated mechanism for Akt inhibition during HCMV reactivation and adds to the growing evidence of the importance of modulating Akt activity in CD34^+^ HPCs.

HCMV reactivation from latency is a multi-step, multi-component process which depends on re-expression of viral genes that are largely silenced during latency, including IE1 and IE2. While IE1 and IE2 transcripts are mostly driven by the MIEP during lytic infection, recent work has uncovered that expression of genes from the MIE locus are driven by two alternative promoters, iP1 and iP2, in CD34^+^ HPCs. Deletion of the transcriptional start sites for iP1 and/or iP2 results in a virus that is unable to reactivate from latency (46). Moreover, FOXO3a binding sites in this region are important for iP1– and iP2-driven transcript expression and reactivation (34). In lytic infection, phosphorylated Akt restricts HCMV replication in a mechanism dependent on inactivating FOXO3a. Expressing a constitutively active Akt impaired IE transcript accumulation, including iP1 and iP2-driven transcripts, resulting in a defect in viral DNA synthesis. However, artificially inducing FOXO3a nuclear localization was able to overcome the inhibition of IE transcription induced by constitutive Akt activity (28). The mechanisms underlying the regulation of Akt and FOXO3a signaling during HCMV infection have not yet been fully elucidated, but our data demonstrate a role for miR-UL36, miR-UL112, and miR-UL148D in reducing Akt expression and signaling and thereby promoting FOXO3a activation and nuclear translocation. Infection of fibroblasts with the ΔmiR-UL36/112/148D virus resulted in reduced iP1– and iP2-driven transcript accumulation (Fig. 6D, E), but no change in MIEP expression (Fig. 6C), supporting a link between Akt levels and iP1 and iP2 transactivation. The ΔmiR-UL36/112/148D mutant is unable to reactivate from latency both *in vitro* and *in vivo*, similar to preventing FOXO3a binding to iP1 and iP2 regions of the viral genome. Critically, introducing an Akt shRNA into the miRNA mutant virus partially restored the ability of the virus to reactivate, underscoring the important role for miRNA-mediated attenuation of Akt signaling during reactivation.

Akt is a central kinase in the cell involved in numerous essential cellular functions, and so reduction of Akt protein levels by HCMV miRNAs may also affect additional processes that contribute to reactivation from latency, such as modulating myeloid differentiation. HCMV infection of monocytes induces Akt activation and drives differentiation into macrophages (71–76), and so further study is needed to assess the effects of reducing Akt signaling on myeloid differentiation. Nevertheless, our findings support previously published work establishing a role for regulation of Akt signaling during infection as well as the requirement of Akt attenuation to allow for FOXO3a activation and IE gene expression, describing for the first time a mechanism employed by HCMV miRNAs to regulate Akt during reactivation.

## MATERIALS AND METHODS

### Cells and media

Feeder-free hESCs were obtained from WiCell (WA01-H1, hPCSC Reg identifier (ID) WAe001-A, NIH approval no. NIHhESC-10-0043). Cells were thawed and plated on Matrigel-coated six-well plates in complete mTeSR1 (Stem Cell Technologies). CD34^+^ HPCs were differentiated using a commercial feeder-free hematopoietic differentiation kit (STEMdiff Hematopoietic Kit, Stem Cell Technologies). HEK293 and adult normal human dermal fibroblasts (NHDF) were obtained from ATCC and cultured in Dulbecco’s modified Eagle’s medium (DMEM) supplemented with 10% heat-inactivated fetal bovine serum (FBS; Hyclone), 100 units/ml penicillin, 100 μg/ml streptomycin, and 100 μg/ml glutamine (Thermofisher). M2-10B4 and S1/S1 stromal cells were obtained from Stem Cell Technologies and maintained in DMEM with 10% FBS and penicillin, streptomycin, and glutamine as previously described (77). All cells were maintained at 37°C and 5% CO2.

### Viruses

Viruses used in this study include BAC-generated WT TB40/E expressing GFP from the SV40 promoter (78, 79). TB40/E mutant viruses containing point mutations in the pre-miRNA sequences for miR-UL36, miR-UL112 and miR-UL148D and viruses expressing either cel-miR-67 or an Akt shRNA in the 3’UTR of UL22A were generated by galactokinase (galK)-mediated recombination (80). Briefly, the galK gene was inserted into the region of the pre-miRNA hairpin using homologous recombination (miR-UL36 galK F: GAAATAAGAAAAATCCACGCACGTTGAAAACACCTGGAAAGAACGTGCCCGAGCGAACGTCCTCTTTCCAGGTGTCCCTGTTGACAATTAATCATCGGCA, miR-UL36 galK R: GCTCCGT TCGCGCAACGCCCTGGGGCCCTTCGTGGGCAAGATGGGCACCGTCTGTTCGCAAGGTAAGCCCCACGCTCAGCAAAAGTTCGATTTATTCAAC, miR-UL12 galK F: CACAGCATGAACACCAGATGCTCCCGGCGCTCTGACAGCCTCCGGATCACATGGTTACTCAGCGTCTGCCAGCCTCCTGTTGACAATTAATCATCGGCA, miR-UL112 galK R: CCTCGGGTTGCCTGGACGCCTGGGCGCGACGCGGCGTGCTGCTGCTCAACACCGTGTTCACCGTGGTGCACGGACTCAGCAAAAGTTCGATTTATTCAAC, miR-UL148D galK F: GAGGCAGAAGCTCGGTTCTCCAGGGACGACCGTCGATGCGTGGTAGGCGCCCTGTTGACAATTAATCAT CGGCA, miR-UL148D galK R: AACTATCTGCAGAACACAAGGAAAAAGAAACACCAACCGAGGGTGGGTGGCTCAGCAAAAGTTCGATTTA), or the galK gene was inserted into the 3’UTR of pUL22A (F: AAGACTGATGAACACAAAGAAAATCAAGCCAAAGAAAATGAAAAGAAGATTCAGTAACAGCAGACCCCAAGGGTTAACGACctgttgacaattaatcatc, R: AAAGAAAAAAGACCGGAGGCGGGGTGTTTTTAGAGCAAAACCTTACAGCTTTTTAATAAAAAACAAGGTAGTCAACATAACTCAGCAAAAGTTCGATTTA). In the second recombination step, galK is removed using oligos that encompass the pre-miRNA sequence containing point mutations in the hairpin (miR-UL36 F: CACCTGGAAAGAACGTGCCCGAGCGAACGTCCTCTTTCCAGGTGTCAAGTTGctCGTGGGGCTTACCTTGCGAACAGACGGTGCCCATCTTGCCCACGAA, miR-UL36 R: TTCGTGGGCAAGATGGGCACCGTCTGTTCGCAAGGTAAGCCCCACGAGCAACTTGACACCTGGAAAGAGGACGTTCGCTCGGGCACGTTCTTTCCAGGTG, miR-UL112 F: AGCCTCCGGATCACATGGTTACTCAGGCTCTGCCAGCCTAAATGCCGGTGAGAGCCCGGCTGTCCGTGCACCACGGTGAACACGGTGTTGAGCAGCAGCA, miR-UL112 R: TGCTGCTGCTCAACACCGTGTTCACCGTGGTGCACGGACAGCCGGGCTCTCACCGGCATTTAGGCTGGCAGACGCTGAGTAACCATGTGATCCGGAGGCT, miR-UL148D F: TGAGGTTGGGGCGGATAACGTGTTGCGGATCGTGGCGAGAACGTGGTGCTACCCTTCTTCACCGCCCCACCCACCCTCGGTTGGTGTTTCTTTTTCCTTG, miR-UL148D R: CAAGGAAAAAGAAACACCAACCGAGGGTGGGTGGGGCGGTGAAGAAGGGTAGCACCACGTTCTCGCCACGATCCGCAACACGTTATCCGCCCCAACCTCA) or inserting a cel-miR-67 (GCTGTTGACAGTGAGCGGCTACTCTTTCTAGGAGGTTGTGATAGTGAAGCCACAGATGTATCACAACCTCCTAGAAAGAGTAGATGCCTACTGCCTCGGA) or Akt shRNA sequence (TGCTGTTGACAGTGAGCGCGCGTGACCATGAACGAGTTTTAGTGAAGCCACAGATGTAAAACTCGTTCATGGTCACGCATGCCTACTGCCTCGGA). All virus stocks were propagated and titered on NHDFs using standard techniques. To assess growth kinetics, NHDFs were infected at a MOI of 3 for single-step growth curves or a MOI of 0.01 for multi-step growth curves for 2 hr. Cell-associated and supernatant virus was harvested at multiple time points post-infection. Titers were determined by plaque assay on NHDFs.

### Reagents

The 3’UTR of human Akt or Pin1 was amplified by PCR from fibroblast genomic DNA and cloned downstream of the *Renilla* luciferase gene in the psiCHECK-2 dual reporter construct (Promega) by XhoI (Akt) or SpeI (Pin1) and NotI restriction sites using the following primer pairs: Akt-GCGGCTCGAGCACACCACCTGACCAAGAT and CGCCGCGGCCGCGAAAAGCAACTTTTATTGAAGAATTTGGAG, Pin1-GCGCACTAGTGCAGAAGCCATTTGAAGACGC and GCGCGCGGCCGCGCAGACAGTGGTTCTGG. siGENOME RISC-free control siRNA (Neg; Dharmacon), Akt siRNA (s659; Thermofisher), and Pin1 siRNA (s10546, Thermofisher) were used in transfection experiments. Double stranded miRNA mimics were custom designed and synthesized by Integrated DNA Technologies. pEGFP-Akt1 (WT) was a gift from Thomas Leonard and Ivan Yudushkin (Addgene plasmid #86637; http://n2t.net/addgene:86637; RRID:Addgene_86637). The following commercial antibodies were used: Akt (C7H310, Cell Signaling), p-Akt S473 (9271, Cell Signaling), p-Akt T308 (13038, Cell Signaling), CREB (48H2, Cell Signaling), p-CREB S133 (87G3, Cell Signaling), FOXO3a (D19A7, Cell Signaling), p-FOXO3a S253 (D18H8, Cell Signaling), GFP (GF28R, Invitrogen), GSK3β (D5C57, Cell Signaling), p-GSK3b S9 (D85E12, Cell Signaling), GAPDH (ab8245, Abcam), HCMV IE2 (MAB810, Sigma Aldrich), mTOR (7C10, Cell Signaling), p-mTOR S2448 (D9C2, Cell Signaling), P70S6K (PA5-17883, Cell Signaling), p-P70S6K T389 (B2H9L2, Thermofisher), Pin1 (3722, Cell Signaling), Phalloidin-AlexaFluor 647 (sc-363797, Santa Cruz Biotechnology), PRAS40 (D23C7, Cell Signaling), p-PRAS40 T246 (D4D2, Cell Signaling), α-rabbit-AlexaFluor 555. Afuresertib was purchased from Selleckchem and BAY1125976 was purchased from MedChemExpress.

### Luciferase assays

HEK293T cells were seeded into 96 well plates and transfected with 100 ng of psiCHECK-2 vector and 100 fmol of negative control or miRNA mimic using Lipofectamine 2000 (Invitrogen). Twenty-four hours after transfection cells were harvested for luciferase assay using the Dual-Glo Reporter Assay Kit (Promega) according to the manufacturer’s instructions. Luminescence was detected using a Veritas microplate luminometer (Turner Biosystems). All experiments were performed in triplicate and presented as mean +/− standard deviation.

### Western blot analysis

Cells were harvested in protein lysis buffer (50mM Tris-HCl pH 8.0, 150mM NaCl, 1% NP40, and protease inhibitors), loading buffer (4X Laemmli Sample Buffer with 2-mercaptoethanol) was added, and lysates were incubated at 95°C for 5 min. Extracts were loaded onto 4-15% acrylamide gels (Biorad), transferred to Immobilon-P membranes (Millipore), and visualized with the specified antibodies. The relative intensity of bands detected by Western blotting was quantified using ImageJ software.

### Quantitative RT-PCR

Reverse transcription-PCR (RT-PCR) was used to quantitate cellular and viral RNA in infected NHDFs. Total RNA was isolated from infected cells using Trizol. cDNA was prepared using 1000ng of total RNA and random hexamer primers. Samples were incubated at 16°C for 30 minutes, 42°C for 30 minutes, and 85°C for 5 minutes. Real-time PCR (Taqman) was used to analyze cDNA levels in infected samples. An ABI StepOnePlus Real Time PCR machine was used with the following program for 40 cycles: 95°C for 15 sec and 60°C for 1 minute. Primer and probe sets for Akt (Hs00178289_m1) and 18S (Hs03928990_g1) were obtained from Thermo Fisher Scientific. Sequence-specific primer pairs for MIEP-, iP1-, and iP2-derived transcripts were used as previously described (46). Relative expression was determined using the 1111Ct method using 18S or GAPDH as the standard control with error bars representing the standard deviation from at least 3 experiments.

### Microscopy

NHDFs were grown on 13mm glass coverslips and infected with WT HCMV, ΔmiR-UL36/112/148D, or mock infected. 72 hpi, coverslips were washed with PBS and fixed with 4% paraformaldehyde in PBS. Cells were permeabilized with 0.25% Triton, blocked with normal goat serum, and stained with the indicated primary antibodies. Coverslips were then washed with PBS containing BSA and 0.1% Triton and incubated with the appropriate fluorophore-conjugated secondary antibodies. Fluorescence was visualized using a LEICA Stellaris 8 microscope using the 63x objective with an NA of 1.4. The fluorophores were excited using 405nm and White Light Lasers. The signals were captured using Leica Stellaris 8 and the Leica Application Suite software. Images were exported as .tiff files and analyzed using ImageJ software.

### CD34^+^ HPC latency and reactivation assays

ifferentiated hESCs were infected with the indicated viruses at an MOI of 2 for 48hr, or were left uninfected, in stem cell media (Iscove’s modified Dulbecco’s medium [IMDM] [Invitrogen] containing 10% BIT serum replacement [Stem Cell Technologies], penicillin/streptomycin, stem cell factor [SCF], FLT3 ligand [FLT3L], interleukin-3 [IL-3], interleukin-6 [IL-6] [all from PeproTech], 50uM 2-mercaptoethanol, and 20ng/ml low-density lipoproteins). Pure populations of viable, infected (GFP^+^) CD34^+^ HPCs were isolated by fluorescence-activated cell sorting (FACS) (BD

FACSAria equipped with 488-, 633-, and 405-nm lasers and running FACSDiva software) and used in latency assays as previously described (77, 78). Briefly, cells were cultured in transwells above irradiated stromal cells (M2-10B4 and S1/S1) for 12 days to establish latency. Virus was reactivated by coculture with NHDF in RPMI medium containing 20% FBS, 1% P/S/G, and 15ng/ml each of G-CSF and GM-CSF in an extreme limiting dilution assay (ELDA). GFP^+^ wells were scored 3 weeks postplating and the frequency of infectious centers was using ELDA software (81). In some experiments, Akt inhibitors 100nM Afuresertib or 50nM BAY1125976 was added to the reactivation culture.

### Engraftment and infection of humanized mice

All animal studies were carried out in strict accordance with the recommendations of the American Association for Accreditation of Laboratory Animal Care. The protocol was approved by the Institutional Animal Care and Use Committee (protocol 0922) at Oregon Health and Science University. NOD-*scid*IL2Rγ_c_^null^ mice were maintained in a pathogen-free facility at Oregon Health and Science University in accordance with procedures approved by the Institutional Animal Care and Use Committee. Both sexes of animals were used. Humanized mice were generated as previously described (49). The animals (12-14 weeks post-engraftment) were treated with 1 ml of 4% Thioglycollate (Brewer’s Media, BD) by intraperitoneal (IP) injection to recruit monocyte/macrophages. At 24hr post-treatment, mice were infected with HCMV TB40/E-infected fibroblasts (approximately 10^5^ PFU of cell-associated virus per mouse) via IP injection. A control group of engrafted mice was mock infected using uninfected fibroblasts. Virus was reactivated as previously described (49).

### Quantitative PCR for viral genomes

DNA from CD34^+^ HPCs was extracted using the two-step TRIZOL (Thermofisher) method according to the manufacturer’s directions. Total DNA was analyzed in triplicate using TaqMan FastAdvanced PCR master mix (Applied Biosystems), and primer and probe for HCMV *UL141* and human β-globin as previously described (82). Copy number was quantified using a standard curve generated from purified HCMV BAC DNA and human β-globin-containing plasmid DNA, and data were normalized assuming two copies of β-globin per cell.

### Statistical analysis

Statistical analysis was performed using GraphPad Prism software (v10) for comparison between groups using student’s t-test, one-way or two-way analysis of variance (ANOVA) with Tukey’s post-hoc test or Bonferroni’s multiple comparison test as indicated. Values are expressed as mean +/− standard deviation or standard error of the mean, as indicated in the figure legends. Significance is highlighted with p<0.05.

## ACKNOWLEDGMENTS

This research utilized the Integrated Pathology Core at the Oregon National Primate Research Center (ONPRC). We are grateful for helpful discussions with Felicia Goodrum, Daniel Streblow, and Andrew Yurochko. We the authors would especially like to acknowledge the mentorship and support of our friend and colleague Jay A. Nelson whose kindness and enthusiasm for science we will carry with us always.

## SUPPLEMENTAL FIGURE LEGENDS

**Figure S1.**
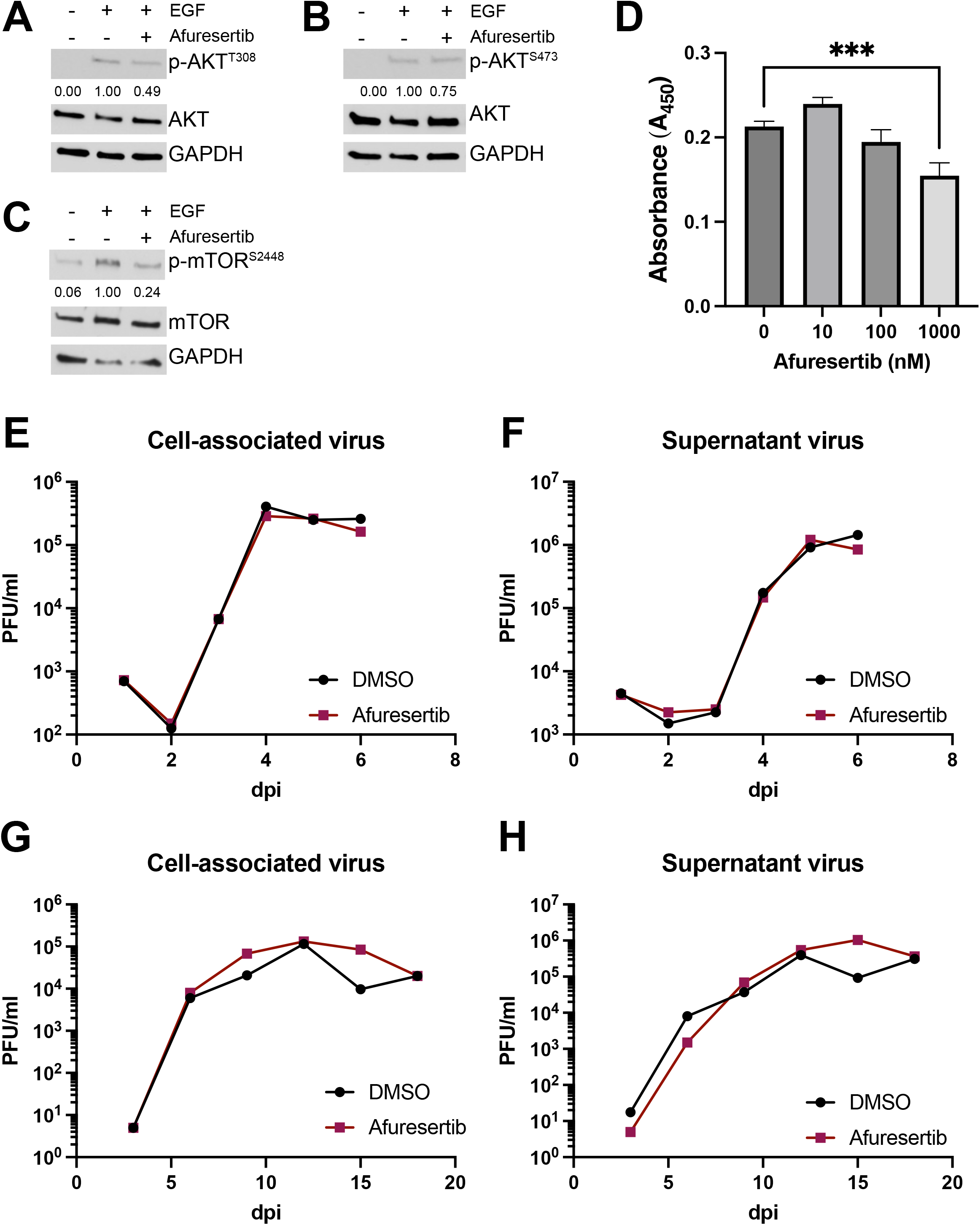
Afuresertib does not affect cell viability or HCMV replication. (A-C) NHDFs were serum starved overnight and treated +/− 100nM Afuresertib. The next day, cells were stimulated +/− EGF for 15 minutes. Protein lysates were harvested and immunoblotted for p-Akt T308 (A), p-Akt S473 (B), or p-mTOR S2448 (C) as well as total Akt (A, B), total mTOR (C), and GAPDH. Quantification from one representative blot shows relative expression levels of p-AKT or p-mTOR compared to cells stimulated with EGF (normalized to GAPDH). (D) CD34+ HPCs were incubated with the indicated concentrations of Afuresertib for 24hr. Cell viability was measured by WST-1 colorimetric assay (Roche) according to manufacturer’s instructions. Quantification shows absorbance at 450nm after background subtracting the value of media alone. Error bars represent standard deviation from triplicate samples (***p<0.0005 [one-way ANOVA with Tukey’s multiple comparison test]). (E-H) NHDFs were infected with WT TB40/E-GFP at an MOI of 3 for single step (E, F) or an MOI of 0.01 for multistep (G, H) growth curves and treated +/− 100nM Afuresertib. PFU/ml values were quantified in duplicate from samples collected at the indicated time points for cell-associated (E, G) or supernatant (F, H) virus.

**Figure S2.**
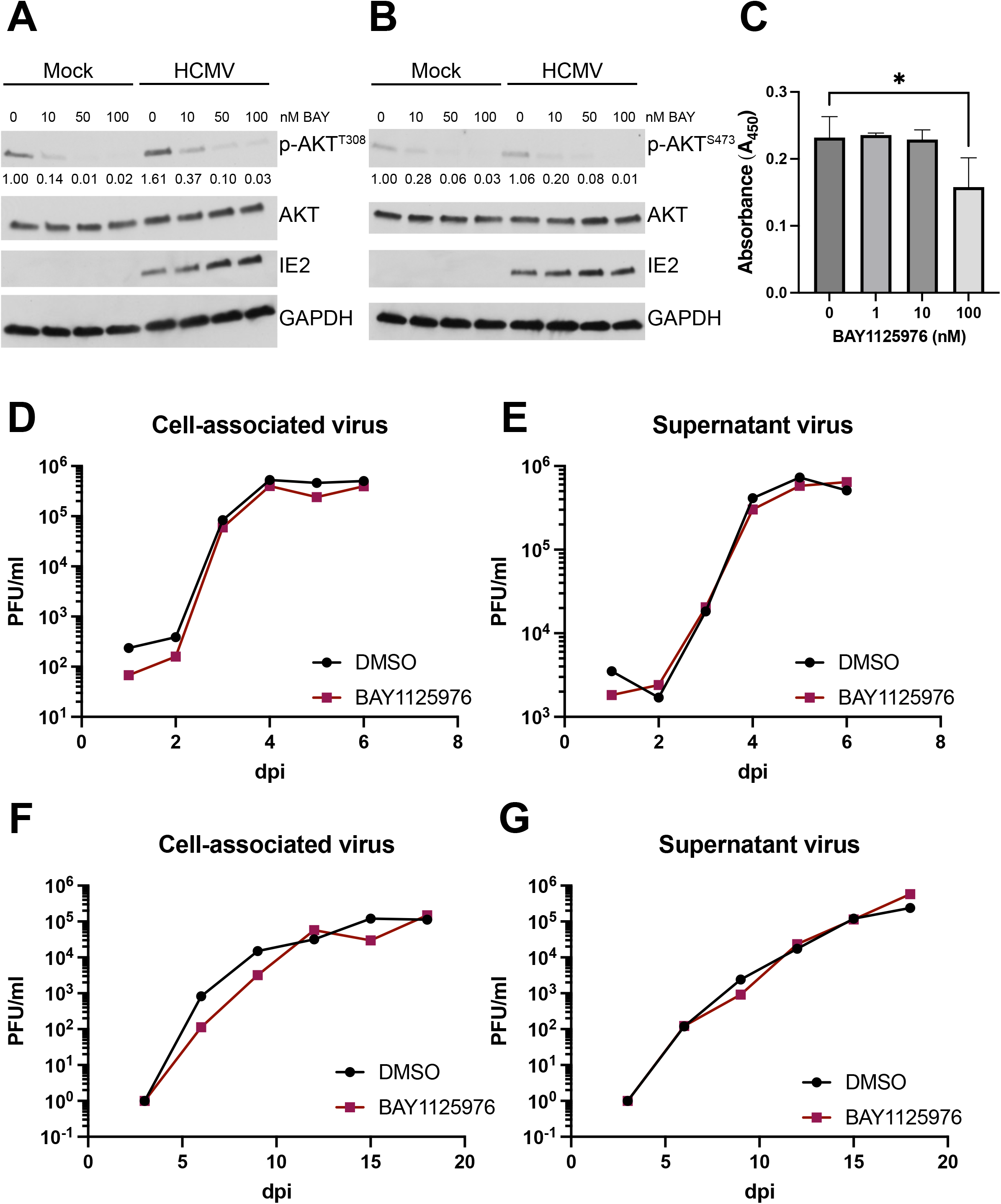
BAY1125976 does not affect cell viability or HCMV replication. (A-C) NHDFs were infected with WT TB40/E-GFP (or Mock infected) at an MOI of 3 for 8hr and then were serum starved overnight in the presence of increasing concentrations of BAY1125976. At 24 hpi, cells were stimulated with EGF for 15 minutes. Lysates were then harvested and immunoblotted for p-Akt T308 (A) or p-Akt S473 (B) as well as total Akt, IE2, and GAPDH. Quantification from one representative blot shows relative expression levels of p-AKT compared to Mock-infected cells stimulated with EGF (normalized to GAPDH). (D) CD34+ HPCs were incubated with the indicated concentrations of BAY1125976 for 24hr. Cell viability was measured by WST-1 colorimetric assay (Roche) according to manufacturer’s instructions. Quantification shows absorbance at 450nm after background subtracting the value of media alone. 27 Error bars represent standard deviation from triplicate samples (*p<0.05 [one-way ANOVA with Tukey’s multiple comparison test]). (E-H) NHDFs were infected with TB40/E-GFP at an MOI of 3 for single-step (E, F) or an MOI of 0.01 for multistep (G, H) growth curves and treated +/− 50nM BAY1125976. PFU/ml values were quantified in duplicate from samples collected at the indicated time points for cell-associated (E, G) or supernatant (F, H) virus.

**Figure S3.**
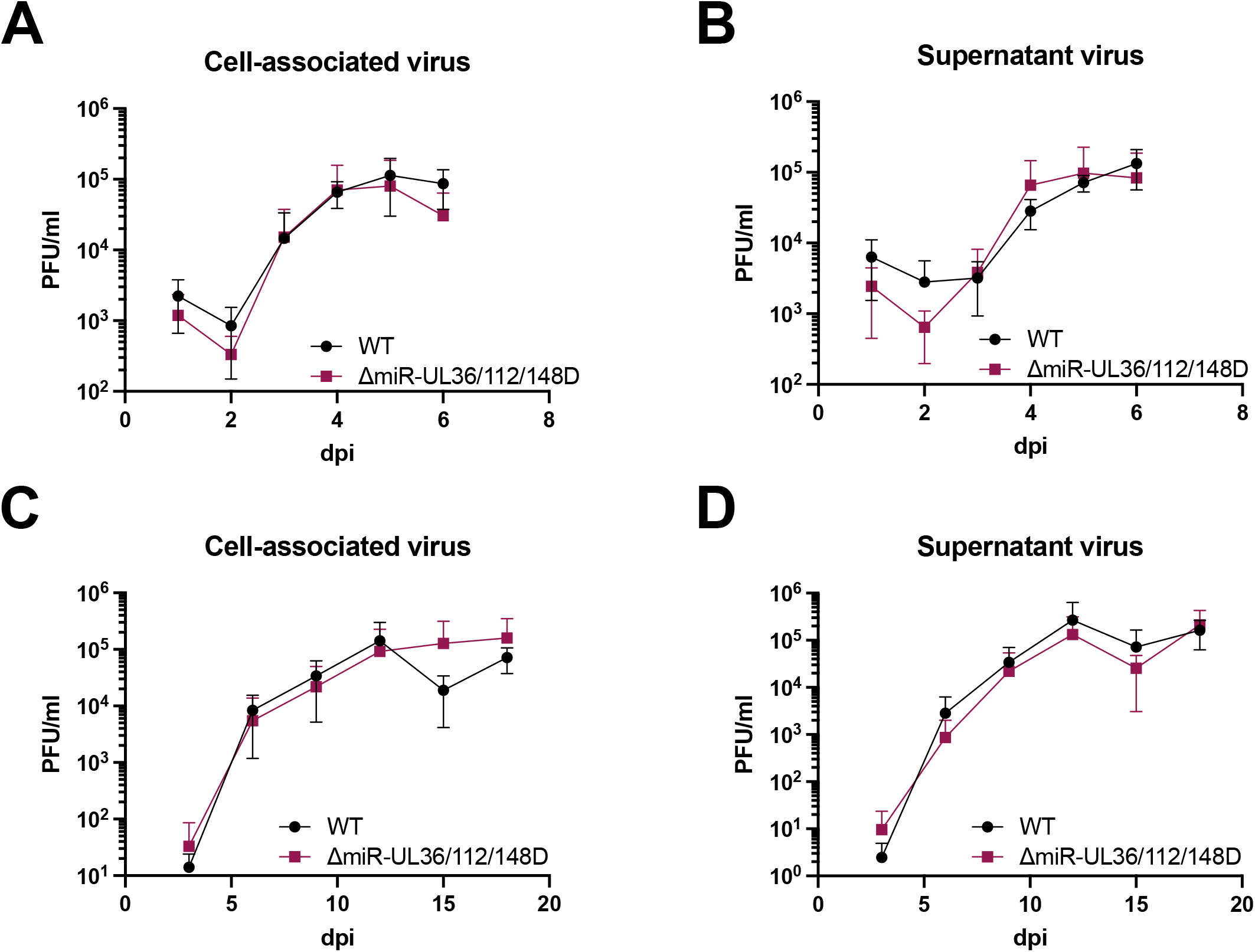
ΔmiR-UL36/112/148D virus does not have a growth defect. NHDFs were infected with WT TB40/E-GFP or ΔmiR-UL36/112/148D at an MOI of 3 for single-step (A, B) or an MOI of 0.01 for multistep (C, D) growth curves. PFU/ml values were quantified in duplicate from samples collected at the indicated time points for cell-associated (A, C) or supernatant (B, D) virus.

**Figure S4.**
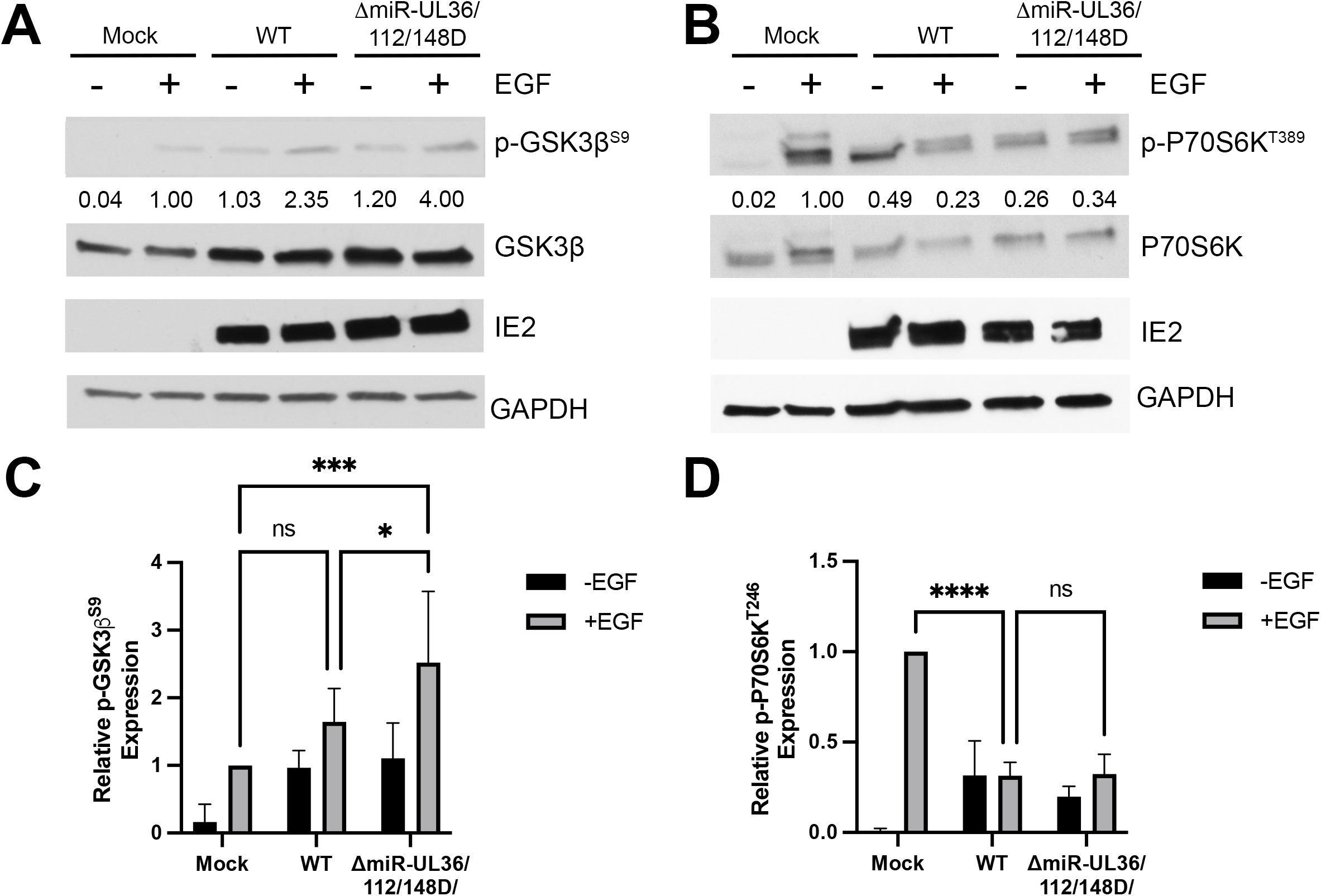
HCMV miR-UL36, miR-UL112, and miR-UL148D alter signaling downstream of Akt. (A, B) NHDF were infected with WT, ΔmiR-UL36/112/148D, or Mock for 48hr, serum starved overnight, and then stimulated +/−EGF for 15 minutes. Lysates were then harvested and immunoblotted for indicated phosphorylated and total proteins as well as HCMV IE2 and GAPDH. Quantification from one representative blot shows relative expression levels of p-protein compared to Neg (normalized to GAPDH). (C, D) Quantification of (A, B), respectively, from three separate experiments (comparing +EGF conditions, *p<0.05, ***p<0.0005, ****p<0.0001 [two-way ANOVA with Tukey’s multiple comparison test]).

**Figure S5.**
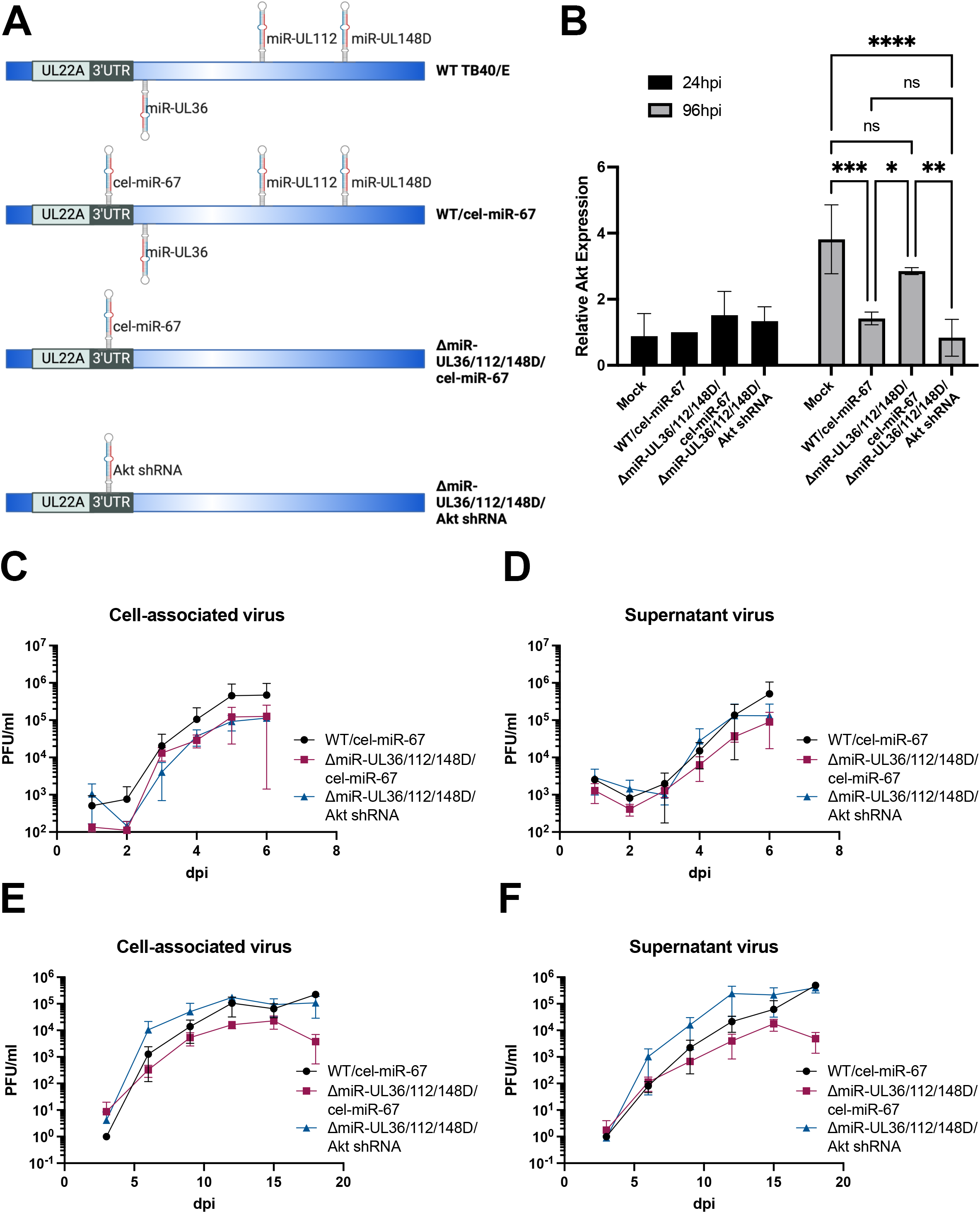
Characterization of the ΔmiR-UL36/112/148D/Akt shRNA virus. (A) Schematic of cel-miR-67 and Akt shRNA-expressing viruses. From top to bottom: WT TB40/E-GFP, WT TB40/E-GFP expressing *C. elegans* miR-67 (cel-miR-67) from the 3’UTR of UL22A (WT/cel-miR-67), ΔmiR-UL36/112/148D expressing cel-miR-67 (ΔmiR-UL36/112/148D/cel-miR-67), or ΔmiR-UL36/112/148D expressing an Akt shRNA from this same region (52 ΔmiR-UL36/112/148D/Akt shRNA) (B) NHDFs were infected with WT/cel-miR-67, ΔmiR-UL36/112/148D/cel-miR-67, ΔmiR UL36/112/148D/Akt shRNA, or Mock for 24 or 96hr after which RNA was harvested. Quantitative RT-PCR was performed using specific primers for Akt. Expression levels were normalized to GAPDH and compared to Mock (p<0.05, **p<0.005, ***p<0.0005, ****p<0.0001 [two-way ANOVA with Tukey’s multiple comparison test]). (C-F) NHDFs were infected with WT/cel-miR-67, ΔmiR UL36/112/148D/cel-miR-67, or ΔmiR-UL36/112/148D/Akt shRNA at an MOI of 3 for single-step (C, D) or an MOI of 0.01 for multistep (E, F) growth curves. PFU/ml values were quantified in duplicate from samples collected at the indicated time points for cell-associated (C, E) or supernatant (D, F) virus.

## REFERENCES

1. Goodrum F. Human Cytomegalovirus Latency: Approaching the Gordian Knot. Annu Rev Virol. 2016;3(1):333–57.

2. Mendelson M, Monard S, Sissons P, Sinclair J. Detection of endogenous human cytomegalovirus in CD34+ bone marrow progenitors. J Gen Virol. 1996;77 (Pt 12):3099–102.

3. Hargett D, Shenk TE. Experimental human cytomegalovirus latency in CD14+ monocytes. Proc Natl Acad Sci U S A. 2010;107(46):20039–44.

4. Ljungman P, Hakki M, Boeckh M. Cytomegalovirus in hematopoietic stem cell transplant recipients. Hematol Oncol Clin North Am. 2011;25(1):151–69.

5. Fulkerson HL, Nogalski MT, Collins-McMillen D, Yurochko AD. Overview of Human Cytomegalovirus Pathogenesis. Methods Mol Biol. 2021;2244:1–18.

6. Salome S, Corrado FR, Mazzarelli LL, Maruotti GM, Capasso L, Blazquez-Gamero D, Raimondi F. Congenital cytomegalovirus infection: the state of the art and future perspectives. Front Pediatr. 2023;11:1276912.

7. Diggins NL, Hancock MH. Viral miRNA regulation of host gene expression. Semin Cell Dev Biol. 2023;146:2–19.

8. Diggins NL, Hancock MH. HCMV miRNA Targets Reveal Important Cellular Pathways for Viral Replication, Latency, and Reactivation. Noncoding RNA. 2018;4(4).

9. Diggins NL, Skalsky RL, Hancock MH. Regulation of Latency and Reactivation by Human Cytomegalovirus miRNAs. Pathogens. 2021;10(2).

10. Diggins NL, Crawford LB, Hancock MH, Mitchell J, Nelson JA. Human Cytomegalovirus miR-US25-1 Targets the GTPase RhoA To Inhibit CD34(+) Hematopoietic Progenitor Cell Proliferation To Maintain the Latent Viral Genome. mBio. 2021;12(2).

11. Hancock MH, Crawford LB, Pham AH, Mitchell J, Struthers HM, Yurochko AD, et al. Human Cytomegalovirus miRNAs Regulate TGF-beta to Mediate Myelosuppression while Maintaining Viral Latency in CD34(+) Hematopoietic Progenitor Cells. Cell Host Microbe. 2020;27(1):104–14 e4.

12. Mikell I, Crawford LB, Hancock MH, Mitchell J, Buehler J, Goodrum F, Nelson JA. HCMV miR-US22 down-regulation of EGR-1 regulates CD34+ hematopoietic progenitor cell proliferation and viral reactivation. PLoS Pathog. 2019;15(11):e1007854.

13. Hancock MH, Mitchell J, Goodrum FD, Nelson JA. Human Cytomegalovirus miR-US5-2 Downregulation of GAB1 Regulates Cellular Proliferation and UL138 Expression through Modulation of Epidermal Growth Factor Receptor Signaling Pathways. mSphere. 2020;5(4).

14. Dunn EF, Connor JH. HijAkt: The PI3K/Akt pathway in virus replication and pathogenesis. Prog Mol Biol Transl Sci. 2012;106:223–50.

15. Liu X, Cohen JI. The role of PI3K/Akt in human herpesvirus infection: From the bench to the bedside. Virology. 2015;479-480:568–77.

16. Chuluunbaatar U, Roller R, Feldman ME, Brown S, Shokat KM, Mohr I. Constitutive mTORC1 activation by a herpesvirus Akt surrogate stimulates mRNA translation and viral replication. Genes Dev. 2010;24(23):2627–39.

17. Soares JA, Leite FG, Andrade LG, Torres AA, De Sousa LP, Barcelos LS, et al. Activation of the PI3K/Akt pathway early during vaccinia and cowpox virus infections is required for both host survival and viral replication. J Virol. 2009;83(13):6883–99.

18. Xiang K, Wang B. Role of the PI3K-AKT-mTOR pathway in hepatitis B virus infection and replication. Mol Med Rep. 2018;17(3):4713–9.

19. Dunn EF, Connor JH. Dominant inhibition of Akt/protein kinase B signaling by the matrix protein of a negative-strand RNA virus. J Virol. 2011;85(1):422–31.

20. Carsillo M, Kim D, Niewiesk S. Role of AKT kinase in measles virus replication. J Virol. 2010;84(4):2180–3.

21. Kumar A, Abbas W, Colin L, Khan KA, Bouchat S, Varin A, et al. Tuning of AKT-pathway by Nef and its blockade by protease inhibitors results in limited recovery in latently HIV infected T-cell line. Sci Rep. 2016;6:24090.

22. Bhaskar PT, Hay N. The two TORCs and Akt. Dev Cell. 2007;12(4):487–502.

23. Kashani B, Zandi Z, Pourbagheri-Sigaroodi A, Yousefi AM, Ghaffari SH, Bashash D. The PI3K signaling pathway; from normal lymphopoiesis to lymphoid malignancies. Expert Rev Anticancer Ther. 2024.

24. Bozulic L, Hemmings BA. PIKKing on PKB: regulation of PKB activity by phosphorylation. Curr Opin Cell Biol. 2009;21(2):256–61.

25. Hers I, Vincent EE, Tavare JM. Akt signalling in health and disease. Cell Signal. 2011;23(10):1515–27.

26. Manning BD, Cantley LC. AKT/PKB signaling: navigating downstream. Cell. 2007;129(7):1261–74.

27. Domma AJ, Henderson LA, Goodrum FD, Moorman NJ, Kamil JP. Human cytomegalovirus attenuates AKT activity by destabilizing insulin receptor substrate proteins. J Virol. 2023;97(10):e0056323.

28. Zhang H, Domma AJ, Goodrum FD, Moorman NJ, Kamil JP. The Akt Forkhead Box O Transcription Factor Axis Regulates Human Cytomegalovirus Replication. mBio. 2022;13(4):e0104222.

29. Bai Y, Xuan B, Liu H, Zhong J, Yu D, Qian Z. Tuberous Sclerosis Complex Protein 2-Independent Activation of mTORC1 by Human Cytomegalovirus pUL38. J Virol. 2015;89(15):7625–35.

30. Moorman NJ, Cristea IM, Terhune SS, Rout MP, Chait BT, Shenk T. Human cytomegalovirus protein UL38 inhibits host cell stress responses by antagonizing the tuberous sclerosis protein complex. Cell Host Microbe. 2008;3(4):253–62.

31. Rodriguez-Sanchez I, Schafer XL, Monaghan M, Munger J. The Human Cytomegalovirus UL38 protein drives mTOR-independent metabolic flux reprogramming by inhibiting TSC2. PLoS Pathog. 2019;15(1):e1007569.

32. Brunet A, Bonni A, Zigmond MJ, Lin MZ, Juo P, Hu LS, et al. Akt promotes cell survival by phosphorylating and inhibiting a Forkhead transcription factor. Cell. 1999;96(6):857–68.

33. Buehler J, Carpenter E, Zeltzer S, Igarashi S, Rak M, Mikell I, et al. Host signaling and EGR1 transcriptional control of human cytomegalovirus replication and latency. PLoS Pathog. 2019;15(11):e1008037.

34. Hale AE, Collins-McMillen D, Lenarcic EM, Igarashi S, Kamil JP, Goodrum F, Moorman NJ. FOXO transcription factors activate alternative major immediate early promoters to induce human cytomegalovirus reactivation. Proc Natl Acad Sci U S A. 2020;117(31):18764–70.

35. Dumble M, Crouthamel MC, Zhang SY, Schaber M, Levy D, Robell K, et al. Discovery of novel AKT inhibitors with enhanced anti-tumor effects in combination with the MEK inhibitor. PLoS One. 2014;9(6):e100880.

36. Yamaji M, Ota A, Wahiduzzaman M, Karnan S, Hyodo T, Konishi H, et al. Novel ATP-competitive Akt inhibitor afuresertib suppresses the proliferation of malignant pleural mesothelioma cells. Cancer Med. 2017;6(11):2646–59.

37. Politz O, Siegel F, Barfacker L, Bomer U, Hagebarth A, Scott WJ, et al. BAY 1125976, a selective allosteric AKT1/2 inhibitor, exhibits high efficacy on AKT signaling-dependent tumor growth in mouse models. Int J Cancer. 2017;140(2):449–59.

38. Medica S, Crawford LB, Denton M, Min CK, Jones TA, Alexander T, et al. Proximity-dependent mapping of the HCMV US28 interactome identifies RhoGEF signaling as a requirement for efficient viral reactivation. PLoS Pathog. 2023;19(10):e1011682.

39. Crawford LB. Human Embryonic Stem Cells as a Model for Hematopoietic Stem Cell Differentiation and Viral Infection. Curr Protoc. 2022;2(12):e622.

40. Hancock MH, Crawford LB, Perez W, Struthers HM, Mitchell J, Caposio P. Human Cytomegalovirus UL7, miR-US5-1, and miR-UL112-3p Inactivation of FOXO3a Protects CD34(+) Hematopoietic Progenitor Cells from Apoptosis. mSphere. 2021;6(1).

41. Bartel DP. Metazoan MicroRNAs. Cell. 2018;173(1):20–51.

42. Liao Y, Wei Y, Zhou X, Yang JY, Dai C, Chen YJ, et al. Peptidyl-prolyl cis/trans isomerase Pin1 is critical for the regulation of PKB/Akt stability and activation phosphorylation. Oncogene. 2009;28(26):2436–45.

43. Kudchodkar SB, Yu Y, Maguire TG, Alwine JC. Human cytomegalovirus infection induces rapamycin-insensitive phosphorylation of downstream effectors of mTOR kinase. J Virol. 2004;78(20):11030–9.

44. Sleman S. Virus-induced FoxO factor facilitates replication of human cytomegalovirus. Arch Virol. 2022;167(1):109–21.

45. Fasano C, Disciglio V, Bertora S, Lepore Signorile M, Simone C. FOXO3a from the Nucleus to the Mitochondria: A Round Trip in Cellular Stress Response. Cells. 2019;8(9).

46. Collins-McMillen D, Rak M, Buehler JC, Igarashi-Hayes S, Kamil JP, Moorman NJ, Goodrum F. Alternative promoters drive human cytomegalovirus reactivation from latency. Proc Natl Acad Sci U S A. 2019;116(35):17492–7.

47. Crawford LB, Diggins NL, Caposio P, Hancock MH. Advances in Model Systems for Human Cytomegalovirus Latency and Reactivation. mBio. 2022;13(1):e0172421.

48. Crawford LB, Streblow DN, Hakki M, Nelson JA, Caposio P. Humanized mouse models of human cytomegalovirus infection. Curr Opin Virol. 2015;13:86–92.

49. Smith MS, Goldman DC, Bailey AS, Pfaffle DL, Kreklywich CN, Spencer DB, et al. Granulocyte-colony stimulating factor reactivates human cytomegalovirus in a latently infected humanized mouse model. Cell Host Microbe. 2010;8(3):284–91.

50. Kim S, Seo D, Kim D, Hong Y, Chang H, Baek D, et al. Temporal Landscape of MicroRNA-Mediated Host-Virus Crosstalk during Productive Human Cytomegalovirus Infection. Cell Host Microbe. 2015;17(6):838–51.

51. Lau B, Poole E, Krishna B, Sellart I, Wills MR, Murphy E, Sinclair J. The Expression of Human Cytomegalovirus MicroRNA MiR-UL148D during Latent Infection in Primary Myeloid Cells Inhibits Activin A-triggered Secretion of IL-6. Sci Rep. 2016;6:31205.

52. Romania P, Cifaldi L, Pignoloni B, Starc N, D’Alicandro V, Melaiu O, et al. Identification of a Genetic Variation in ERAP1 Aminopeptidase that Prevents Human Cytomegalovirus miR-UL112-5p-Mediated Immunoevasion. Cell Rep. 2017;20(4):846–53.

53. Pan C, Zhu D, Wang Y, Li L, Li D, Liu F, et al. Human Cytomegalovirus miR-UL148D Facilitates Latent Viral Infection by Targeting Host Cell Immediate Early Response Gene 5. PLoS Pathog. 2016;12(11):e1006007.

54. Wang YP, Qi Y, Huang YJ, Qi ML, Ma YP, He R, et al. Identification of immediate early gene X-1 as a cellular target gene of hcmv-mir-UL148D. Int J Mol Med. 2013;31(4):959–66.

55. Hancock MH, Hook LM, Mitchell J, Nelson JA. Human Cytomegalovirus MicroRNAs miR-US5-1 and miR-UL112-3p Block Proinflammatory Cytokine Production in Response to NF-kappaB-Activating Factors through Direct Downregulation of IKKalpha and IKKbeta. mBio. 2017;8(2).

56. Stern-Ginossar N, Saleh N, Goldberg MD, Prichard M, Wolf DG, Mandelboim O. Analysis of human cytomegalovirus-encoded microRNA activity during infection. J Virol. 2009;83(20):10684–93.

57. Kim Y, Lee S, Kim S, Kim D, Ahn JH, Ahn K. Human cytomegalovirus clinical strain-specific microRNA miR-UL148D targets the human chemokine RANTES during infection. PLoS Pathog. 2012;8(3):e1002577.

58. Landais I, Pelton C, Streblow D, DeFilippis V, McWeeney S, Nelson JA. Human Cytomegalovirus miR-UL112-3p Targets TLR2 and Modulates the TLR2/IRAK1/NFkappaB Signaling Pathway. PLoS Pathog. 2015;11(5):e1004881.

59. Hook LM, Grey F, Grabski R, Tirabassi R, Doyle T, Hancock M, et al. Cytomegalovirus miRNAs target secretory pathway genes to facilitate formation of the virion assembly compartment and reduce cytokine secretion. Cell Host Microbe. 2014;15(3):363–73.

60. Grey F, Meyers H, White EA, Spector DH, Nelson J. A human cytomegalovirus-encoded microRNA regulates expression of multiple viral genes involved in replication. PLoS Pathog. 2007;3(11):e163.

61. Murphy E, Vanicek J, Robins H, Shenk T, Levine AJ. Suppression of immediate-early viral gene expression by herpesvirus-coded microRNAs: implications for latency. Proc Natl Acad Sci U S A. 2008;105(14):5453–8.

62. Sapkota S, Pillman KA, Dredge BK, Liu D, Bracken JM, Kachooei SA, et al. On the rules of engagement for microRNAs targeting protein coding regions. Nucleic Acids Res. 2023;51(18):9938–51.

63. Zhang K, Zhang X, Cai Z, Zhou J, Cao R, Zhao Y, et al. A novel class of microRNA-recognition elements that function only within open reading frames. Nat Struct Mol Biol. 2018;25(11):1019–27.

64. Hausser J, Syed AP, Bilen B, Zavolan M. Analysis of CDS-located miRNA target sites suggests that they can effectively inhibit translation. Genome Res. 2013;23(4):604–15.

65. Elcheva I, Goswami S, Noubissi FK, Spiegelman VS. CRD-BP protects the coding region of betaTrCP1 mRNA from miR-183-mediated degradation. Mol Cell. 2009;35(2):240–6.

66. Dangelmaier C, Manne BK, Liverani E, Jin J, Bray P, Kunapuli SP. PDK1 selectively phosphorylates Thr(308) on Akt and contributes to human platelet functional responses. Thromb Haemost. 2014;111(3):508–17.

67. Sarbassov DD, Guertin DA, Ali SM, Sabatini DM. Phosphorylation and regulation of Akt/PKB by the rictor-mTOR complex. Science. 2005;307(5712):1098–101.

68. Krishna BA, Wass AB, Dooley AL, O’Connor CM. CMV-encoded GPCR pUL33 activates CREB and facilitates its recruitment to the MIE locus for efficient viral reactivation. J Cell Sci. 2021;134(5).

69. Grey F, Tirabassi R, Meyers H, Wu G, McWeeney S, Hook L, Nelson JA. A viral microRNA down-regulates multiple cell cycle genes through mRNA 5’UTRs. PLoS Pathog. 2010;6(6):e1000967.

70. Lau B, Poole E, Van Damme E, Bunkens L, Sowash M, King H, et al. Human cytomegalovirus miR-UL112-1 promotes the down-regulation of viral immediate early-gene expression during latency to prevent T-cell recognition of latently infected cells. J Gen Virol. 2016;97(9):2387–98.

71. Chan G, Bivins-Smith ER, Smith MS, Smith PM, Yurochko AD. Transcriptome analysis reveals human cytomegalovirus reprograms monocyte differentiation toward an M1 macrophage. J Immunol. 2008;181(1):698–711.

72. Chan G, Nogalski MT, Yurochko AD. Activation of EGFR on monocytes is required for human cytomegalovirus entry and mediates cellular motility. Proc Natl Acad Sci U S A. 2009;106(52):22369–74.

73. Cojohari O, Peppenelli MA, Chan GC. Human Cytomegalovirus Induces an Atypical Activation of Akt To Stimulate the Survival of Short-Lived Monocytes. J Virol. 2016;90(14):6443–52.

74. Smith MS, Bentz GL, Alexander JS, Yurochko AD. Human cytomegalovirus induces monocyte differentiation and migration as a strategy for dissemination and persistence. J Virol. 2004;78(9):4444–53.

75. Smith MS, Bentz GL, Smith PM, Bivins ER, Yurochko AD. HCMV activates PI(3)K in monocytes and promotes monocyte motility and transendothelial migration in a PI(3)K-dependent manner. J Leukoc Biol. 2004;76(1):65–76.

76. Smith MS, Bivins-Smith ER, Tilley AM, Bentz GL, Chan G, Minard J, Yurochko AD. Roles of phosphatidylinositol 3-kinase and NF-kappaB in human cytomegalovirus-mediated monocyte diapedesis and adhesion: strategy for viral persistence. J Virol. 2007;81(14):7683–94.

77. Crawford LB, Hancock MH, Struthers HM, Streblow DN, Yurochko AD, Caposio P, et al. CD34(+) Hematopoietic Progenitor Cell Subsets Exhibit Differential Ability To Maintain Human Cytomegalovirus Latency and Persistence. J Virol. 2021;95(3).

78. Umashankar M, Goodrum F. Hematopoietic long-term culture (hLTC) for human cytomegalovirus latency and reactivation. Methods Mol Biol. 2014;1119:99–112.

79. Sinzger C, Hahn G, Digel M, Katona R, Sampaio KL, Messerle M, et al. Cloning and sequencing of a highly productive, endotheliotropic virus strain derived from human cytomegalovirus TB40/E. J Gen Virol. 2008;89(Pt 2):359–68.

80. Warming S, Costantino N, Court DL, Jenkins NA, Copeland NG. Simple and highly efficient BAC recombineering using galK selection. Nucleic Acids Res. 2005;33(4):e36.

81. Hu Y, Smyth GK. ELDA: extreme limiting dilution analysis for comparing depleted and enriched populations in stem cell and other assays. J Immunol Methods. 2009;347(1-2):70–8.

82. Crawford LB, Kim JH, Collins-McMillen D, Lee BJ, Landais I, Held C, et al. Human Cytomegalovirus Encodes a Novel FLT3 Receptor Ligand Necessary for Hematopoietic Cell Differentiation and Viral Reactivation. mBio. 2018;9(2).

